# Early Immune Hypoactivation and Persistent Innate Reprogramming Characterize Chronic Chikungunya Disease

**DOI:** 10.64898/2026.06.22.733701

**Authors:** Lien De Caluwé, Candice Bohaud, Borita Heng, Sokchea Lay, Sopheak Sorn, Sreymom Ken, Alvino Maestri, Veasna Duong, Sowath Ly, Giorgio Gonnella, Tineke Cantaert

## Abstract

Chikungunya virus (CHIKV) infection causes acute febrile illness and debilitating arthralgia, with up to 40-80% of patients developing chronic chikungunya disease characterized by persistent arthralgia and fatigue lasting months to years. The immunological mechanisms underlying this transition remain poorly understood.

We performed longitudinal immune profiling of CHIKV-infected patients stratified by clinical outcome. Single-cell RNA sequencing of peripheral blood mononuclear cells collected during acute infection and six months post-infection was combined with multiplex plasma cytokine and flow cytometric analysis.

Patients who later developed chronic symptoms were characterized by early innate immune hypoactivation during acute infection, characterized by reduced monocyte and dendritic cell frequencies, lower circulating IFN-α, IL-6 and IL-8, and reduced antigen-presentation and interferon-associated transcriptional programs. This was accompanied by diminished adaptive immune activation, including reduced IL-17 and IL-21 and lower *HLA-DR* expression across multiple T-cell subsets. Cell–cell communication analysis further indicated impaired acute monocyte-centered immune coordination in chronic progressors, whereas non-chronic patients displayed stronger monocyte-driven innate-to-adaptive signaling. By six months, chronic patients showed persistent innate remodeling, including increased non-classical monocytes, altered plasmacytoid dendritic cell and natural killer cell transcriptional programs, elevated CXCL10, reduced IL-6 and MMP8, and emergence of NK-centered predicted communication networks. Longitudinal transcriptomic analysis further identified divergent immune recovery trajectories, most prominently in non-classical monocytes and plasmacytoid dendritic cells.

Together, these findings suggest that chronic chikungunya disease is associated with early innate immune hypoactivation followed by persistent innate immune remodeling, providing insight into immune mechanisms that may contribute to post-viral chronic inflammatory syndromes.

## Introduction

Chikungunya virus (CHIKV), an alphavirus from the *Togaviridae* family, was first isolated in 1952 in present-day Tanzania. Due to the stooped posture and rigid gait that infected individuals show, the disease was named “chikungunya” derived from the Makonde root verb *kungunyala* meaning “that which bends up” [1–3]. CHIKV is mainly transmitted by the bite of infected female *Aedes* mosquitoes, primarily *Aedes aegypti* and *Aedes albopictus.* It belongs to a group of arthritogenic alphaviruses that includes Ross River virus, o’nyong’nyong virus, and Mayaro virus. CHIKV is the most thoroughly investigated arthritogenic alphavirus, especially regarding its potential to cause long-term chronic illness [2–4].

Chikungunya fever is an acute febrile disease characterized by high fever, myalgia, joint pain, headache, rash, and high viremia, lasting for several days. During the acute phase, the high viremia in the blood allows CHIKV to be passed on from an infected person to its mosquito vector. The onset of disease follows an average incubation period of 3 days, but this can vary between 2 to 12 days [1, 3, 5]. Most infected individuals are symptomatic, only around 15% of them have asymptomatic seroconversion [1]. Joint pains associated with CHIKV infection are usually symmetric and localized in both arms and legs and in both the large and small joints [1, 3, 5]. Approximately 40-80% of chikungunya patients develop chronic chikungunya disease which is characterized by sustained, debilitating joint pain that lasts several months or even years and poses a significant economic and societal burden [3, 6–8].

Despite causing significant morbidity with long-term impact on patient quality of life, no specific therapies for chronic chikungunya exist. The main features of chronic chikungunya include myalgia and arthralgia, closely resembling rheumatoid arthritis [3, 9]. Some known risk factors associated with chronic disease include high viral load, older age and female sex [10–12].

While the exact causes of chronic chikungunya disease remain unclear, two main pathogenic mechanisms have been suggested. The first one is the presence of persistent viral infection or viral antigens in the joints. Long-term viral persistence in macrophages has been described in nonhuman primates and mouse models [13, 14]. Persistent CHIKV was identified in synovial macrophages of a chronic patient from La Réunion, however, similar findings were not observed in synovial fluid samples from chronic patients in Colombia [10, 15]. Alternatively, chronic chikungunya may result from sustained, abnormal immune responses involving macrophage and T cell infiltration in the joints. Studies comparing immune responses between chronic and non-chronic chikungunya patients are limited and focus on the analysis of serum cytokines. The acute phase of CHIKV infection is characterized by elevated levels of inflammatory mediators such as CXCL10, IL-6, IFN-α and CCL2 compared with either convalescent phase or healthy donors, whereas increased levels of IL-6, IL-17 and GM-CSF have been associated with the development of chronic chikungunya disease [3, 16–19]. CD4L T cells have been repeatedly implicated in the immunopathogenesis of CHIKV-mediated arthritis. Experimental and clinical studies have shown that CD4L T cells infiltrate infected joint tissues and promote inflammatory responses that contribute to joint swelling and disease severity, particularly during the acute phase of infection [10, 20, 21].

Despite its high prevalence, there are no biomarkers or mechanistic models that reliably predict which patients will progress to chronic disease. To address this gap, we established a prospective longitudinal cohort during the 2020 CHIKV outbreak in Cambodia. CHIKV-positive individuals were sampled during acute infection and again six months later, at which time they were stratified according to clinical outcome. Using single-cell RNA (scRNA) sequencing of peripheral blood mononuclear cells (PBMCs) and multiplex cytokine profiling, we sought to characterize the cellular and molecular immune trajectories that distinguish chronic from non-chronic cases both in the acute and chronic phase of the disease.

## Materials and Methods

### CHIKV patient recruitment

Participant enrolment was initiated by Institut Pasteur du Cambodge (IPC) in response to the CHIKV epidemic in Cambodia in 2020-2021. Ethical approval for the study was obtained from the National Ethics Committee of Health Research of Cambodia (NECHR #2020-312). Written informed consent was obtained from all participants or parents/legal guardians prior to inclusion in the study. Individuals who tested positive for CHIKV by RT-qPCR during the acute phase of infection, were included in the study [22]. At the six months timepoint participants were clinically examined by the study nurse for presence or absence of persistent arthralgia symptoms (joint swelling, pain or redness). Participants with persistent symptoms in at least one joint were classified as chronic chikungunya cases, whereas patients with absence of joint symptoms were classified as non-chronic [7, 23].

At inclusion, EDTA blood tubes were collected while at the six months timepoint both EDTA and heparin blood tubes were collected. After collection, blood was centrifuged and plasma collected. PBMCs were isolated from EDTA blood using Ficoll–Paque (GE Healthcare) density gradient centrifugation. PBMCs were cryopreserved in liquid nitrogen, and plasma samples were stored at -80 °C until analysis. At the six months timepoint only heparin plasma samples were stored.

For scRNAseq and flow cytometry studies, patients were selected based on sample availability and sample quality (scRNAseq: PBMC viability <70%; flow cytometry: PBMC viability of <25 % were discarded). Patient groups were matched for age and sex as much as possible (**Table 1).**

**Table 1:**
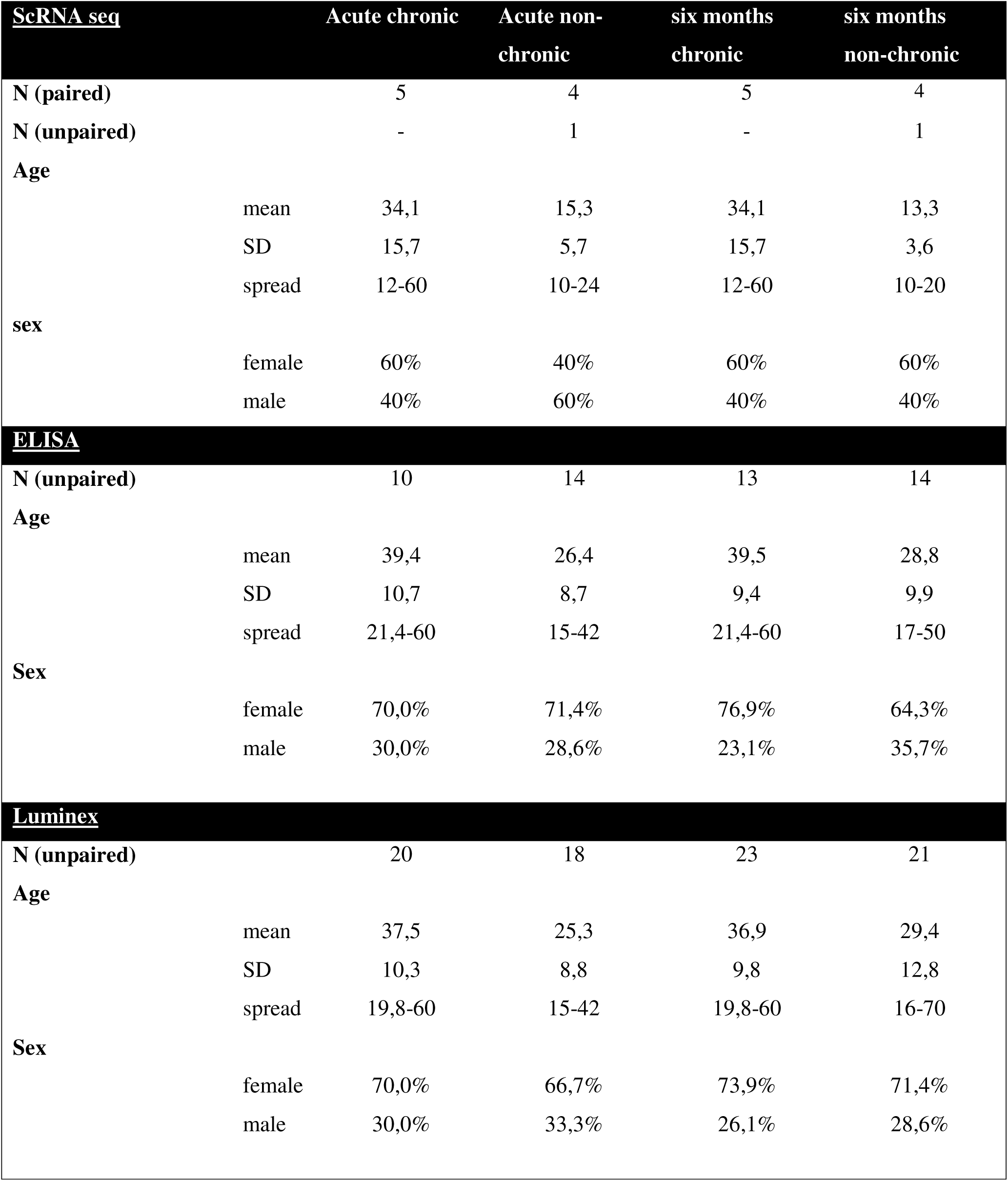
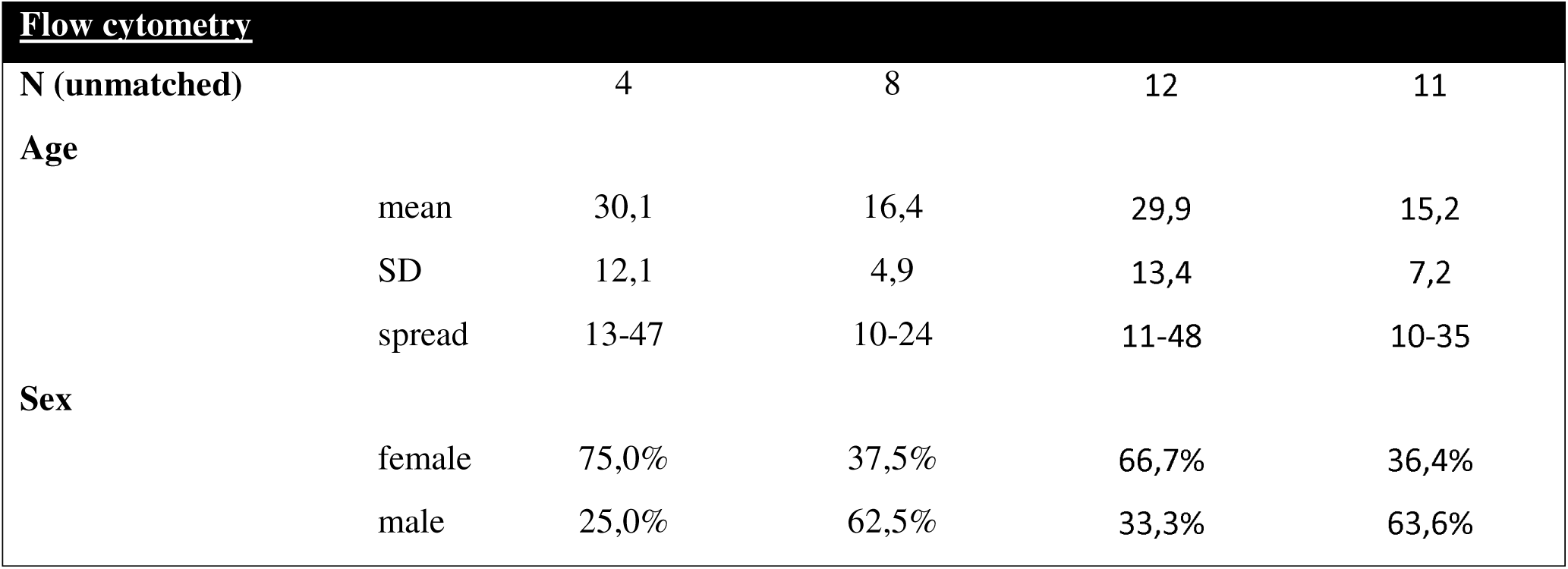
patient and sample characteristics.

### Multiplex Luminex assay

Unpaired acute EDTA plasma samples and six months heparin plasma samples were used to determine analyte concentrations **(Table 1**). A Luminex Human discovery assay (R&D systems) custom-designed to contain 23 analytes was used. For analysis only comparisons within the acute samples and within the six months samples were performed. Samples were processed according to the manufacturer’s instructions. All samples were analyzed in duplicate and were diluted 2-fold. Samples from the different groups were equally distributed over the multiple plates. All standard curves were fitted using a 5-parameter logistic (5PL) model unless the fit was interrupted or yielded inadequate results due to lack of convergence, interpolation failure, or better performance in extrapolating low or high concentration values. In these cases, a 4-parameter logistic (4PL) model was applied instead. Values below the predefined detection limit according to the manufacturer’s instructions or out of range values were excluded from the analysis.

### TGF-β1 ELISA

On the same sample set, TGF-β1 levels were analyzed using the Quantikine ELISA Human/Mouse/Rat/Porcine/Canine TGF-β1 (R&D systems) in unmatched acute EDTA plasma samples and six months heparin plasma samples (**Table 1**). Plasma samples were thawed and centrifuged 10,000 g for 10 min whereafter samples were activated using sample activation kit 1 (R&D systems DY010). Activated samples were diluted 12-fold to 30-fold and processed according to the manufacturer’s instructions. Standard curve was constructed by plotting absorbance of each standard and generating a 4PL curve.

### Flow cytometry

Unpaired PBMCs samples were selected for flow cytometric analysis (**Table 1**). Frozen cells were quickly thawed in a 37 °C warm water bath and subsequently transferred to pre-warmed (37 °C) Iscove’s Modified Dulbecco’s medium (IMDM; Gibco) with 10 % FBS. Cells were washed with PBS and Fc receptors on the PBMCs were blocked with Human TruStain FcX Fc Receptor Blocking Solution (BioLegend). Subsequently cells were stained for chemokine receptors listed in Supplementary Table 1 for 30min at 37 °C. Hereafter the cells were washed with PBS and stained for 20 min at 4 °C with Zombie Aqua fixable viability kit (Biolegend) to determine live/dead cells. Cells were washed with PBS containing 0.5 % BSA and 2 mM EDTA and stained for cell membrane markers for 30 min at 4 °C. After membrane staining, cells were fixed and permeabilized using True-Nuclear Transcription Factor Buffer Set (BioLegend) per manufacturer’s instruction followed by intracellular staining of IκBα (**Supplementary Table 1**). The cells were analyzed with BD FACSAria Fusion (BD Biosciences). Subpopulations were determined using FlowJo (**Supplementary Table 1 and supplementary figure 1**), and mean fluorescence intensity was calculated. In each experiment, single-stained controls for BV421 (IκBα), APC (TGFBR2), and BV785 (CD69) were used to normalize MFI across runs. Samples were excluded if they had less than 25 % viable cells or had less than 12,000 viable cells. The minimum viable cell threshold ensured sufficient cell numbers for reliable analysis. Mean fluorescence intensity was only calculated if the gated population had more than 50 events.

### scRNA sequencing

Longitudinal paired PBMC patients (n=5/per group) (except one unpaired patient in the non-chronic group) were selected for scRNA sequencing analysis (**Table 1**). Cells were stained for 20 min at 4 °C with Zombie Aqua fixable viability kit (BioLegend) to determine live/dead cells. Cells were washed with PBS containing 0.5 % BSA (Sigma-Aldrich) and 2 mM EDTA (Sigma-Aldrich) whereafter the viable cells were sorted using BD FACSAria Fusion (BD Biosciences) into PBS supplemented with 0.2 % BSA. Following cell sorting, 5’ gene expression (GEX) libraries were prepared using the Chromium Next GEM Single Cell 5’ Reagent Kit v2 (10X Genomics), according to the manufacturer’s protocol. Briefly, 15,000 cells per sample were loaded onto the Chromium Controller (10X Genomics) to generate single-cell gel beads-in-emulsion (GEMs). Following reverse transcription, GEMs were broken, and barcoded cDNA was recovered and amplified by PCR. Amplified cDNA was fragmented, end-repaired, and A-tailed before adapter ligation. Sample indexes were added during index PCR, and purified libraries were obtained after size selection. Sequencing was performed by Macrogen (Seoul, Korea) on an Illumina HiSeq X Ten platform using 150 bp paired-end reads. Targeted sequencing depths were ∼20,000 reads per cell for GEX libraries.

### scRNA data processing

Raw sequencing data were processed using Cell Ranger (10x Genomics) v.7.1.0 [24] to generate gene–barcode count matrices aligned to the GRCh38 reference genome. Downstream analysis was performed in R (v.4.5.3) based on the Seurat package (v.5.1.0) [25]. The analysis workflow employed rmdreportdeck v.0.2.1 (https://github.com/baia-ipc/rmdreportdeck) for interactive report generation. Preprocessing was implemented in the baia.seurat.helpers library v.0.1.1 (https://github.com/baia-ipc/baia.seurat.helpers). Thereby cells were filtered using sample-specific cutoffs for the number of detected genes and UMIs and mitochondrial read fraction, derived from scater (v.1.32.1) [26]. Additionally, we eliminated cells identified as upper-tail outliers by robustbase (v.0.99-2) [27] outlyingness scores. Gene expression counts were then processed with Seurat, including normalization, variable-feature selection, scaling, PCA, Harmony integration of different samples (v1.2.3) [28], and Uniform Manifold Approximation and Projection (UMAP) [29] embedding.

Cell types were assigned by SingleR (v.2.1.0) [30] using the MonacoImmuneData reference dataset [31] at fine resolution. Cell type proportions were compared using scProportionTest (v0.0.0.9000) [32]. Differential gene expression analysis was performed on pseudo-bulked counts using DESeq2 (v.1.48.1) [33], as implemented in the baia.pseudobulk.deseq v.0.1.1 (https://github.com/baia-ipc/baia.pseudobulk.deseq) library. Thereby, prior to analysis, MALAT1, mitochondrial, TCR, immunoglobulin, and ribosomal protein genes were removed and significant results were defined by adjusted p-value < 0.05. Gene set enrichment analyses (GSEA) were implemented in the baia.pathway.analysis v.0.1.1 (https://github.com/baia-ipc/baia.pathway.analysis) library, based on clusterProfiler (v.4.16.0) and fgsea (v.1.34.0), using the Reactome database, obtained through the ReactomePA package (v.1.52.0).

The results were reduced into broader biological themes to minimize redundancy among overlapping pathways. Themes were manually curated based on pathway nomenclature and function, and pathways were assigned using keyword matching. For each theme and cell type, the pathway with the highest absolute normalized enrichment score (NES) was selected as representative for visualization.

### Longitudinal scRNA sequencing data analysis

For each immune cell type, longitudinal transcriptional changes between acute infection and six months post infection were quantified separately for chronic and non-chronic patient groups. Log2 fold changes (log_2_FC) for the acute → six months comparison were defined as:

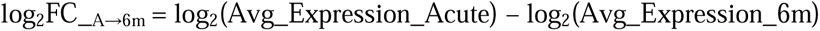

Under this definition, positive values indicate higher expression during the acute phase (declining toward six months), whereas negative values indicate higher expression at six months (increasing during recovery). In addition, differential expression between chronic and non-chronic patients at either acute or six months was quantified as:

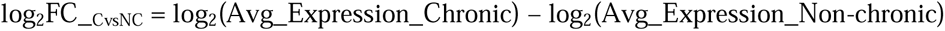

with positive values indicating higher expression in chronic patients and negative values indicating higher expression in non-chronic patients.

To quantify whether longitudinal transcriptional trajectories differed between clinical outcomes, we computed a Δlog statistic defined as:

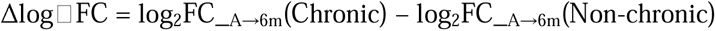

This statistic represents a difference-in-differences contrast, comparing the within-group longitudinal expression change (acute → six months) between chronic and non-chronic patients. ΔlogJFC values quantify differences in longitudinal expression trajectories between chronic and non-chronic patients. ΔlogJFC was used as an effect-size metric to rank genes showing divergent longitudinal trajectories rather than as a formal statistical test. ΔlogJFC values were calculated for genes significantly differentially expressed in at least one of the two longitudinal comparisons (chronic or non-chronic acute → 6 months). For pathway analysis, we further restricted to genes with |ΔlogJFC | ≥ 1, focusing on substantial differences in recovery trajectories **(Supplementary Figure 2)**.

Over-representation analysis (ORA) was performed per cell type using fora() from the fgsea (v 1.36.2) R package, with Reactome gene sets restricted to pathways of 5–500 genes after intersection with the tested-gene universe. Pathogen-specific and nervous-system Reactome terms were excluded. Multiple testing correction used the Benjamini–Hochberg procedure (FDR ≤ 0.05). Redundant pathways were collapsed using a Jaccard similarity threshold of 0.60. A shared gene set was additionally defined as genes with |Δlog| ≥ 1 in at least six immune cell types and subjected to the same ORA framework.

Thus, ΔlogJFC was used as an effect-size metric to quantify differences in longitudinal recovery trajectories between clinical groups, while statistical significance was assessed at the gene level by differential expression testing and at the pathway level by ORA with FDR correction.

### CellChat data analysis

Cell–cell communication networks were inferred using the CellChat framework, as implemented in the baia.cellchat v.0.2.1 helper library, which wraps CellChat v2.2.0 and follows the analytical workflow of Jin et al. [34]. Normalized RNA expression values and cell-type annotations from the Seurat object were used as input. For each condition, CellChat objects were constructed and annotated with the human CellChatDB database, and processed by identifying overexpressed genes and interactions, smoothing expression over the human protein–protein interaction network, computing communication probabilities, filtering low-support interactions, inferring pathway-level communication probabilities, aggregating interaction networks, and estimating pathway centrality. Condition-specific CellChat objects were then merged for pairwise comparisons between acute non-chronic and chronic, and between six months non-chronic and chronic samples. Global communication differences were visualized using interaction-count and interaction-weight heatmaps and differential interaction plots. Signaling roles were summarized for both outgoing and incoming patterns using signaling-role heatmaps. Pathway-level activity was assessed with rankNet-based comparisons of relative and absolute information flow, enabling identification of changes in signaling strength, signaling pathways, and sender/receiver cell populations across conditions.

### Statistical analysis

Flow cytometry data were analyzed using FlowJo version 10.8.1. Statistical analyses for flow cytometry, ELISA, and multiplex cytokine data were performed using GraphPad Prism 9.0 (GraphPad Software, USA). Data are presented as median with interquartile range (IQR). For unpaired data, the Mann–Whitney U test was used to compare two groups. A p-value < 0.05 was considered statistically significant. Multiple testing correction for transcriptomic analyses was performed as described in the scRNA sequencing data analysis section. All statistical tests were two-sided.

## Results

### Longitudinal CHIKV cohort for immune profiling

To assess the contribution of the immune response to chronic arthralgic disease outcome after CHIKV infection, we performed in-depth immunological profiling during the acute phase of infection and at six months post-infection. This included scRNA sequencing, exploratory immunophenotyping in a limited set of patients, and plasma cytokine analysis at both timepoints.

We established a prospective longitudinal cohort during the 2020 CHIKV outbreak in Cambodia (**Figure 1A**). Patients with laboratory confirmed CHIKV infection and who were sampled within 96h of onset of fever were included [22]. Patients were followed up at six months, patients were evaluated for persistent joint pain. Patients were classified as a chronic chikungunya case in case at least 1 joint showed persistent symptoms (swelling, pain, redness) [7, 23]. Chronic patients reported persistent arthralgia affecting both large and small peripheral joints, most frequently the knees, ankles, wrists, and small joints of the hands and feet (**Figure 1A**).

**Figure 1:**
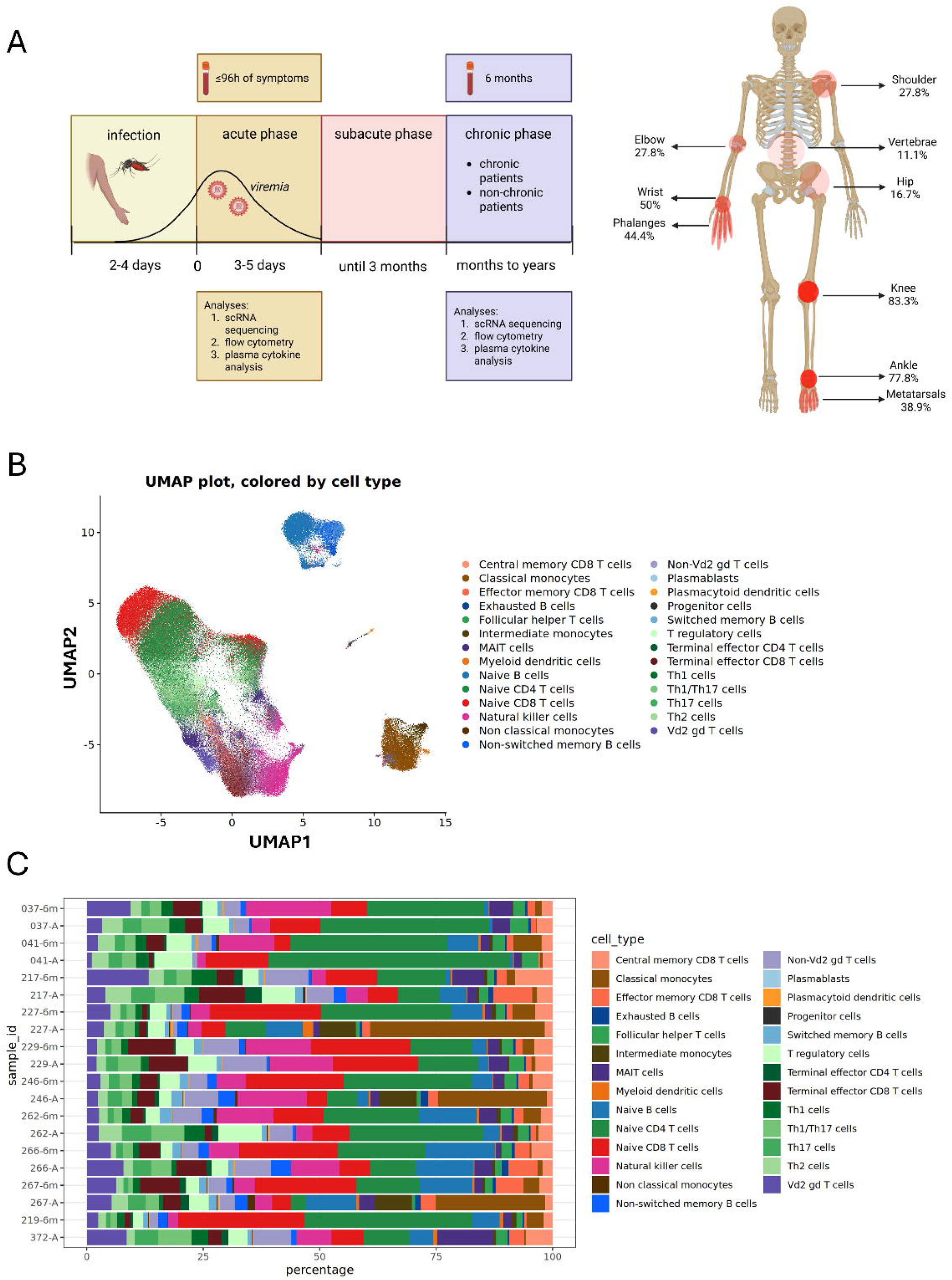
Study design and integrated single-cell atlas of CHIKV infection. **(A)** Graphical representation of the study design. RT-qPCR CHIKV-positive patients were enrolled during the acute phase of infection (≤96 hours after symptom onset). Participants were followed and reassessed at six months post-infection, when they were stratified into chronic and non-chronic outcome groups based on the presence or absence of persistent arthralgia. Anatomical distribution of joint pain reported at the six months follow-up among chronic patients. Created in Biorender. **(B)** UMAP visualization of integrated PBMCs single-cell transcriptomes after Harmony correction, colored by annotated immune cell populations. **(C)** Relative abundance of annotated immune cell populations across individual samples shown as stacked bar plots.

Single cell RNAsequencing was performed in acute and paired 6-month samples of 5 self-resolving and 5 chronic chikungunya patients. Following quality control, filtering, and batch correction using Harmony, PBMCs from all samples were integrated into a shared low-dimensional embedding and visualized by UMAP. Cells were annotated into major immune populations based on canonical marker expression (**Figure 1B**). The integrated dataset showed the expected PBMCs organization, with lymphoid populations (T and natural killer (NK) cells) forming a continuous compartment clearly separated from myeloid populations (monocytes and myeloid dendritic cells (mDCs)), while B-cell subsets formed a distinct cluster. Less abundant populations, including plasmacytoid dendritic cells (pDCs) and progenitor cells, localized together in a separate compartment.

This overall structure was consistent across individual samples, with no evident sample- or batch-specific clustering, supporting robust data integration and reliable cell type annotation (**Supplementary Figure 3**). Although the global immune landscape was preserved across clinical outcome groups and timepoints, inter-individual variation in cell type composition was observed, particularly in the relative abundance of major lymphoid and myeloid populations (**Figure 1C**). These data established a harmonized single-cell atlas for subsequent analyses of immune correlates associated with chronic arthralgia after CHIKV infection.

### Early innate immune dampening characterizes acute infection in patients who develop chronic CHIKV disease

We hypothesized that differences in disease outcome are already dictated at the earliest phase of disease, during the acute phase of infection. Therefore, we first evaluated differences in gene expression, immune cell phenotype and cytokine production in acute CHIKV disease patients who later developed chronic disease compared to patients with non-chronic chikungunya. scRNA sequencing analysis revealed a lower relative abundance of several innate immune populations in patients who later developed chronic disease, including classical, intermediate, and non-classical monocytes, as well as mDCs and pDCs (**Figure 2A).** These differences were not recapitulated by exploratory flow cytometry, likely reflecting technical differences between platforms and the limited flow cytometry sample size **(Supplementary Figure 4**).

**Figure 2:**
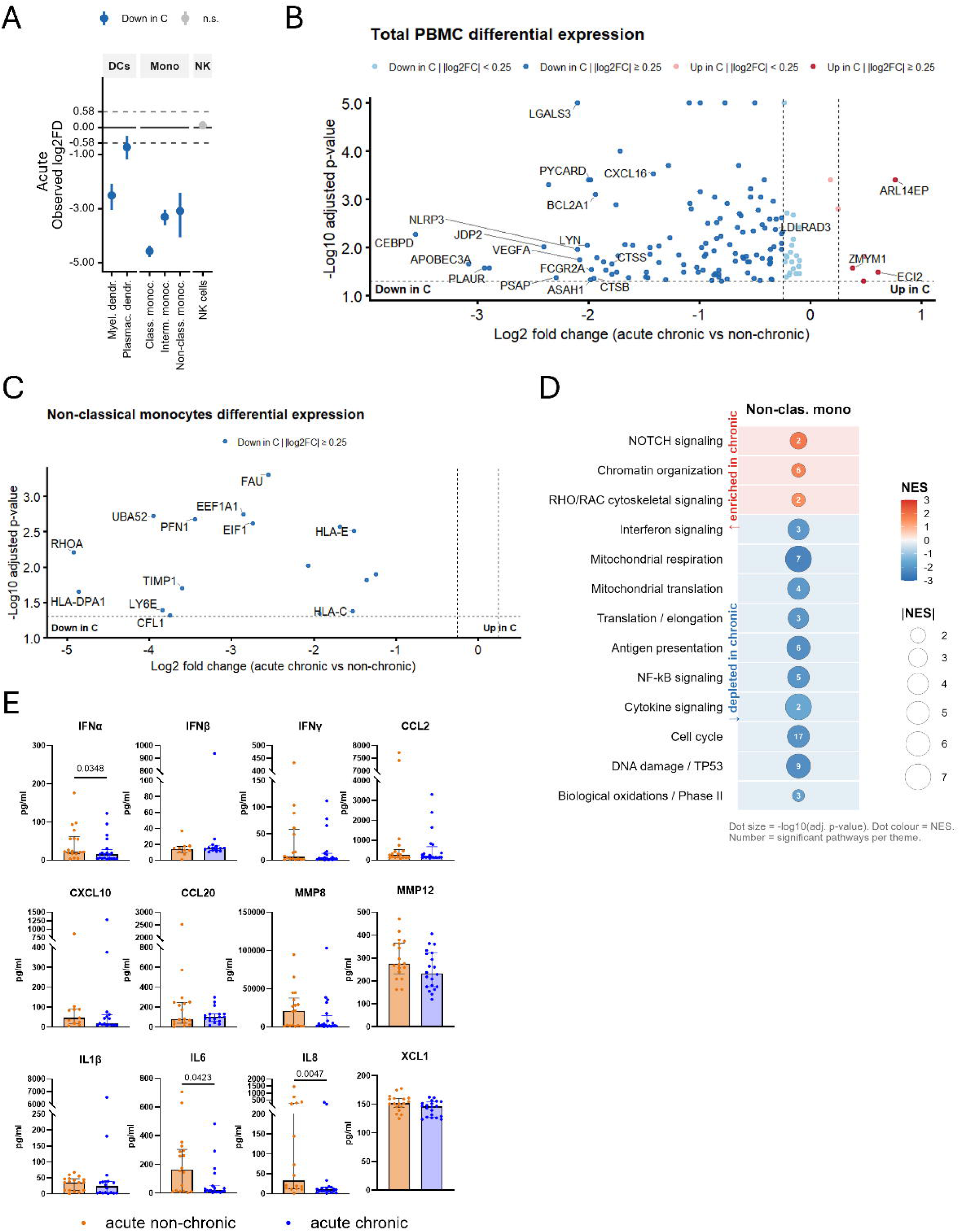
Early attenuation of monocyte-driven innate immunity in chronic CHIKV patients during the acute phase of infection. **(A)** Cell-type proportion differences obtained from the acute phase scRNA sequencing analysis. Log2 fold-differences (FD) represent abundance differences between chronic (n=5) and non-chronic (n=5) patients, where positive values indicate higher frequencies in chronic individuals and negative values indicate higher frequencies in non-chronic individuals. Significant changes are highlighted in blue (|log□FD| ≥ 0.58; FDR < 0.05). Boxplots show the median and interquartile range, with individual points representing each sample. **(B-C)** Differential gene expression in total PBMCs and non-classical monocytes during acute infection. Negative log□ fold-change values indicate downregulation in chronic patients. The volcano plot highlights genes significantly altered between groups (adjusted p-value < 0.05). **(D)** Reactome GSEA analysis of non-classical monocytes during acute infection. Related Reactome pathways were consolidated into broader functional themes for visualization. Red indicates increased normalized enrichment score (NES) in chronic patients, blue indicates depleted enrichment zone in chronic patients. Dot size represents the absolute |NES| of the top-ranked pathway within each theme. Dot fill indicates pathway significance (−log10 adjusted p-value). Numbers within dots indicate the number of significantly enriched pathways assigned to each theme. Red-shaded rows indicate themes relatively enriched in chronic patients; blue-shaded rows indicate themes relatively depleted in chronic patients. **(E)** Acute phase plasma cytokine and chemokine concentrations in non-chronic (orange) and chronic (blue) patients (n = 10–23 per analyte, depending on assay performance). Bars represent median with interquartile range; dots represent individual patients. Exact two-sided Mann–Whitney tests were used for group comparisons.

In total PBMCs, 145 significant differentially expressed genes (DEGs) were identified, of which the majority were downregulated in chronic patients. Chronic patients exhibited coordinated downregulation of genes involved in inflammasome activation (*NLRP3, PYCARD*), transcriptional regulation of inflammatory responses (*CEBPD, JDP2*), myeloid cell identity and phagocytic capacity (*LYN, FCGR2A, BCL2A1*), lysosomal and antigen processing (*CTSB, CTSS, ASAH1, PSAP*), tissue remodeling (*VEGFA, LGALS3, PLAUR*), and the antiviral restriction factor *APOBEC3A*. Only a limited number of genes were upregulated in chronic patients during the acute phase. Among these, *ARL14EP*, a regulator of intracellular vesicular trafficking linked to antigen presentation pathways, showed the strongest effect. Interestingly, downregulation of *LDLRAD3*, a known entry receptor for Venezuelan equine encephalitis virus, another alphavirus, but not CHIKV, was observed (**Figure 2B**) [35].

Among individual immune subsets, non-classical monocytes showed the strongest acute transcriptional divergence between outcome groups, with the highest number of DEGs across annotated cell populations (**Supplementary Figure 5A).** In chronic patients, these cells exhibited coordinated downregulation of genes involved in cytoskeletal organization (*RHOA, PFN1, ACTB, CFL1*), antigen processing and presentation (*HLA-DPA1, HLA-C, HLA-E*), translation machinery (*UBA52, EEF1A1, FAU, EIF1*), interferon-stimulated antiviral response (*LY6E*), and matrix/cytokine regulation (*TIMP1*). The broad suppression of both cytoskeletal and translational genes in non-classical monocytes is noteworthy, as these processes are tightly coupled to monocyte motility, phagocytosis, and effector cytokine production (**Figure 2C**). Consistent with this, Reactome GSEA showed depletion of cytokine signaling, interferon signaling, NF-κB signaling, antigen presentation, mitochondrial respiration, translation-related pathways, and RHO downstream effector signaling including ROCK, Citron kinase, and Rhotekin/Rhophilin pathways, in non-classical monocytes from chronic patients. In contrast, NOTCH signaling, chromatin organization, and RHO/RAC GTPase cycling were relatively enriched in chronic patients, suggesting a potential imbalance between upstream RHO/RAC GTPase cycling and downstream cytoskeletal effector programs **(Figure 2D)**.

This cellular dampening was mirrored at the systemic level: chronic patients showed significantly lower circulating concentrations of key innate cytokines and chemokines, including IFN-α, IL-6, and IL-8, with a trend towards lower concentrations for XCL1 (Hodges–Lehmann median difference -8.0 pg/mL, 95% CI −19.3 to 0.6; *p* = 0.066) and MMP12 (Hodges–Lehmann median difference -48.0 pg/mL, 95% CI −110.0 to 10.2; *p* = 0.091) (**Figure 2E**, **Supplementary Figure 6**).

Together, these findings indicate that patients who progressed to chronic chikungunya displayed an attenuated acute innate immune response, characterized by reduced circulating innate immune cell abundance, broad transcriptional suppression in non-classical monocytes, and lower systemic inflammatory mediator production.

### Acute adaptive immune activation is blunted in chronic patients

Next, we aimed to understand the earliest differences in adaptive immune cell responses during the acute phase of CHIKV infection in patients with non-chronic or chronic disease. scRNA sequencing revealed increased frequencies of T follicular helper (Tfh), naïve CD4J T and naïve CD8J T cells, together with reduced frequencies of CD4J terminal effector (TEMRA) cells, CD8J effector memory (TEM) cells and γδ T cells in patients developing chronic disease. These effector/memory populations are generally associated with cytotoxic antiviral responses, suggesting reduced differentiation toward effector T cell states during acute infection (**Figure 3A**). These findings could not be confirmed by flow cytometry, likely due to the limited sample size and differences in subset resolution between platforms (**Supplementary Figure 4**).

**Figure 3:**
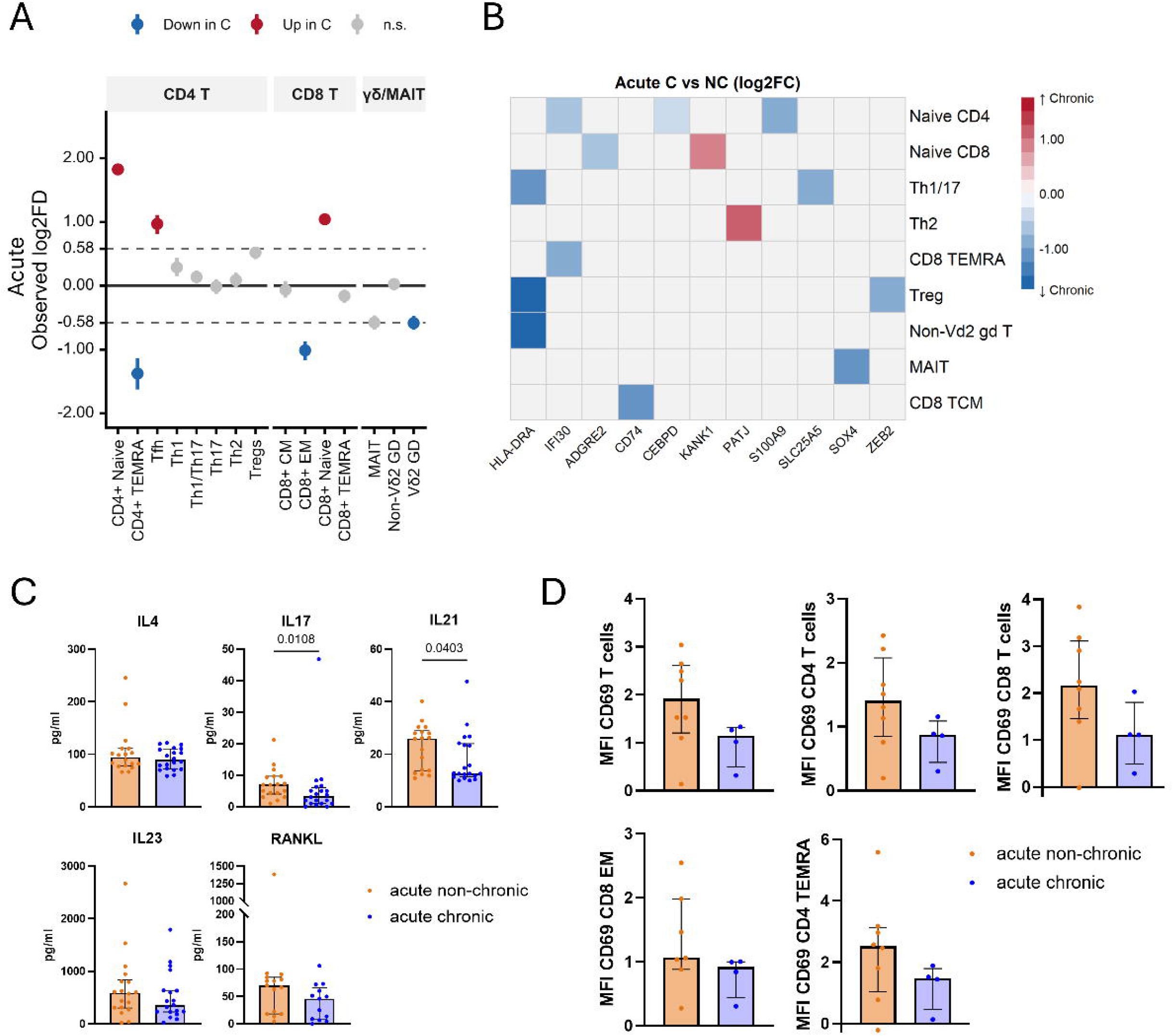
Acute adaptive immune activation is diminished in chronic CHIKV patients. **(A)** Cell-type proportion differences obtained from the acute phase scRNA sequencing analysis. Log fold-differences (FD) represent abundance differences between chronic and non-chronic patients, where positive values indicate higher frequencies in chronic individuals and negative values indicate higher frequencies in non-chronic individuals. Significant changes are highlighted in red and blue (|log□FD| ≥ 0.58; FDR < 0.05). Boxplots show the median and interquartile range, with individual points representing each sample. **(B)** Differential gene expression in adaptive immune subsets at the acute timepoint. Heatmap summarizes significantly DEGs between non-chronic and chronic patients across key T- and B-cell subsets. Negative log□ fold-change values indicate downregulation in chronic patients. **(C)** Acute phase plasma cytokine concentrations associated with T cell activation and differentiation (n = 13–23 per analyte, depending on assay performance). Bar plots show individual donor values, median with IQR is shown. Statistical comparisons performed using the Mann–Whitney U test. **(D)** CD69 expression levels on CD4□ and CD8□ T cells during acute infection, quantified as mean fluorescence intensity (MFI) by flow cytometry (n = 4–8). Bars represent median with interquartile range; dots represent individual patients. Exact two-sided Mann–Whitney tests were used for group comparisons.

Transcriptional profiling revealed impaired activation across adaptive immune subsets in chronic chikungunya patients. Notably, *HLA-DR* was consistently downregulated in multiple T cell populations, including Th1/Th17 cells, regulatory T cells, and non-Vδ2 γδ T cells, suggesting a widespread reduction in T cell activation (**Figure 3B**).

At the systemic level, patients developing chronic disease had significantly lower concentrations of IL-17 and IL-21, with trends toward reduced RANKL (Hodges–Lehmann median difference -18.4 pg/mL, 95% CI −59.0 to 3.1; *p* = 0.077) compared to patients with non-chronic disease (**Figure 3C and Supplementary Figure 6)**. These reductions reflect diminished T helper activity and impaired coordination of both humoral and cellular responses.

Flow cytometry provided partial support for these early adaptive differences. T cells from chronic patients showed a trend (Hodges–Lehmann median difference -1.1, 95% CI −2.0 to 0.18; *p* = 0.1091) toward reduced expression of CD69, an early activation marker, indicating reduced early T cell activation (**Figure 3D**).

Together, these cellular, cytokine, and transcriptional findings converge on a coherent picture of early adaptive hyporesponsiveness in chronic patients with reduced frequencies of cytotoxic T cell subsets, reduced T cell activation and impaired cytokine production. This weakened initiation of effector responses could drive the divergent immune recovery trajectories observed later in infection.

### Distinct innate and adaptive immune alterations characterize chronic chikungunya at six months post-infection

Chronic chikungunya is defined by a persistence of clinical symptoms 3 to 6 months post-infection [3]. To better understand the immunological alterations associated with chronic disease, we profiled both the innate and adaptive immune compartments six months post-infection using scRNA sequencing, immunophenotyping, and plasma cytokine analysis.

scRNA sequencing revealed that the frequency of non-classical monocytes was higher in chronic chikungunya patients, indicating altered monocyte skewing in chronic disease (**Figure 4A**). Together with this altered myeloid profile, chronic patients also displayed higher frequencies of NK cells and lower frequencies of naïve CD8J T cells and CD4^+^ TEMRA cells (**Figure 4A**).

**Figure 4:**
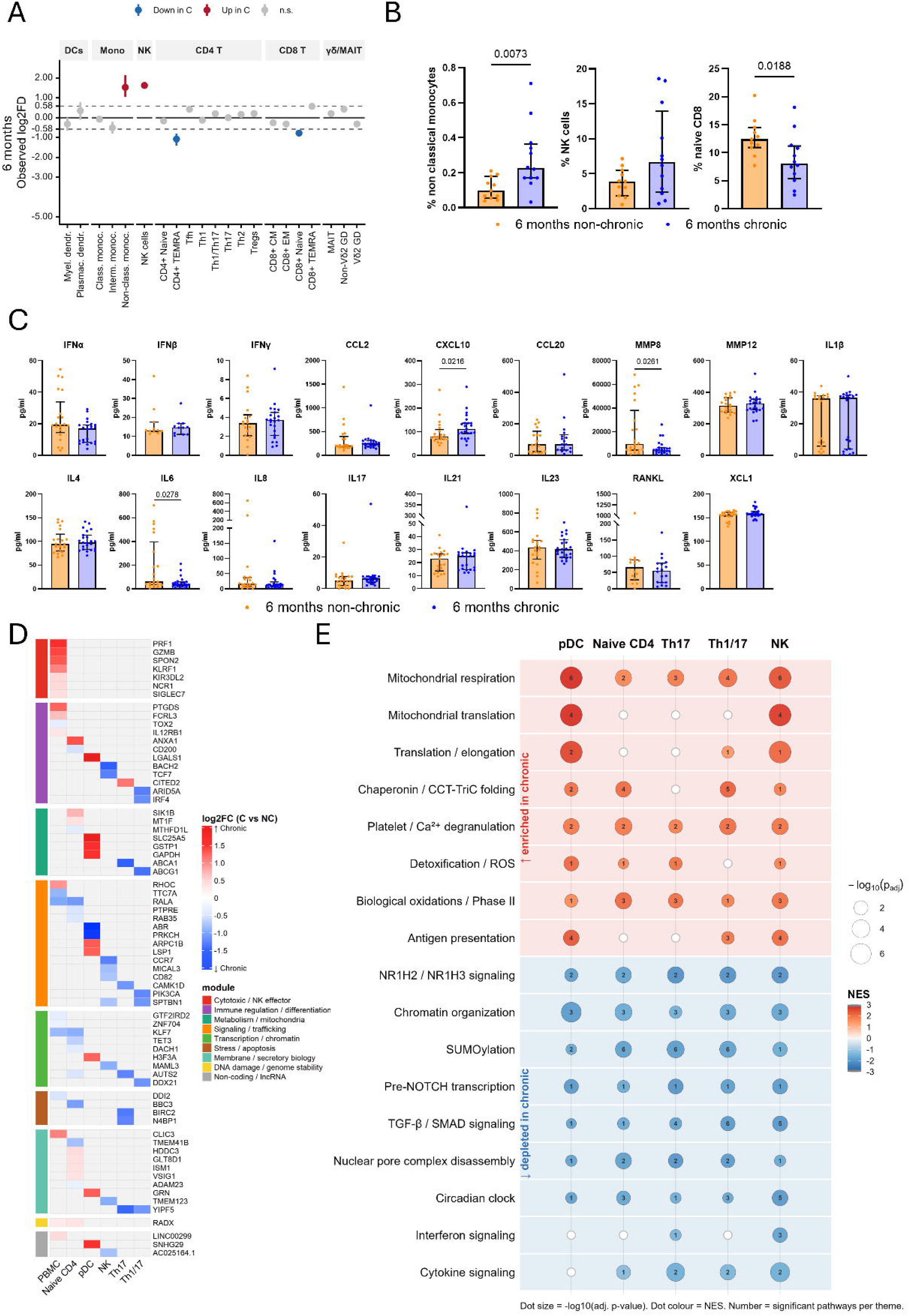
Distinct immune composition, cytokine profile, and transcriptional programs characterize chronic chikungunya at six months post-infection. **(A)** Cell-type proportion differences obtained from six months scRNA sequencing analysis. Log fold-differences (FD) represent abundance differences between chronic (n=5) and non-chronic (n=5) patients, where positive values indicate higher frequencies in chronic individuals and negative values indicate higher frequencies in non-chronic individuals. Significant changes are highlighted in red or blue (|log□FD| ≥ 0.58; FDR < 0.05). Plots show the median and interquartile range, with individual points representing each sample. **(B)** Flow-cytometry–derived frequencies of cell subsets among total viable cells, shown as bar plots (n = 11-12). Cells were defined as non-classical monocytes (CD3□/CD14^dim^/CD16□), Th17 cells (CD3□/CD4□/CD45RA□/CCR4□/CCR6□/CXCR3□), NK cells (CD3^−^/CD56^+^). CD4□ and CD8□ T cell subsets were defined as naïve (CD45RA^+^/CCR7^+^), TEMRA (CD45RA□/CCR7□), gated on CD3□CD4□ or CD3□CD8□ T cells respectively. Each point represents an individual donor. Comparisons between chronic and non-chronic patients were performed using the Mann–Whitney U test **(C)** Six months plasma cytokine and chemokine concentrations (n = 10–23 per analyte, depending on assay performance). Bar plots display individual donor values. Statistical comparisons between chronic and non-chronic groups were performed using the Mann–Whitney U test. **(D)** Heatmap of DEGs across PBMCs and immune subsets. DEGs were defined as |log_2_FC| > 0.25 (chronic vs non-chronic) and grouped into co-regulated functional modules. Color scale indicates log2 fold change, with red representing higher expression in chronic patients and blue representing lower expression. **(E)** Pathway enrichment analysis across immune subsets. Differentially enriched pathways were grouped into higher-order functional themes based on shared biological processes. Dot color represents NES, with red indicating enrichment in chronic patients and blue indicating depletion. Dot size reflects statistical significance (−log10 adjusted p-value). Numbers indicate the number of significant pathways per functional theme. Only the top-ranked pathway per theme is shown for visualization.

Overall immunophenotyping further confirmed these alterations, with significant higher frequencies in non-classical monocytes (Hodges–Lehmann median difference 0.13, 95% CI 0.04 to 0.27; p = 0.0073), significant lower frequencies of naïve CD8^+^ T cells (Hodges–Lehmann median difference -4.28, 95% CI −7.79 to -0.90; p = 0.0188) and a trend towards higher frequencies of NK cells (Hodges–Lehmann median difference 2.785, 95% CI −0.86 to 8.51; p = 0.1693) (**Figure 4B).**

At the systemic level, CXCL10 concentrations were elevated in chronic patients, while IL-6 and MMP8 were lower compared to non-chronic patients at six months post infection (**Figure 4C**). IFN-α concentrations showed a decreasing trend in chronic patients that did not reach significance (Hodges–Lehmann median difference −6.19, 95% CI −12.3 to 0.11; p = 0.0572), mirroring the pattern observed in the acute phase. Th associated cytokines, including IL-21 and IL-17, did not differ significantly between groups at six months, indicating that the acute phase reduction in Th-derived cytokines did not persist over time (**Figure 4C, Supplementary Figure 6**).

At the transcriptional level, DEGs were identified across total PBMCs and individual immune subsets, with the highest numbers observed in naïve CD4J T cells, pDCs, NK cells, Th17, and Th1/Th17 cells

(**Supplementary Figure 5B**). DEGs were filtered at |log_2_FC| > 0.25 and grouped into co-regulated modules revealing coordinated transcriptional programs across cell types (**Figure 4D**).

A consistent opposing pattern in directionality was observed: pDCs and naïve CD4J T cells showed a large group of upregulated genes in chronic patients, whereas NK, Th17, and Th1/Th17 cells showed predominantly downregulated gene expression (**Supplementary Figure 5B**), suggesting divergent immune trajectories.

In total PBMCs, upregulated genes clustered within a cytotoxic/NK effector module including *PRF1*, *GZMB*, *SPON2*, *KLRF1*, *KIR3DL2*, and *NCR1*. This signature was not mirrored by cytotoxic gene upregulation within CD8J T-cell subsets, which showed only few DEGs and no canonical cytotoxic genes (**Supplementary Figure 5C**). In contrast, NK cells showed a larger transcriptional divergence, together with increased NK cell frequency in chronic patients. These observations suggest that the PBMC-level cytotoxic signature is most likely NK-associated rather than CD8J T-cell–driven. Together with elevated CXCL10 levels, this points to a sustained innate cytotoxic tone at the systemic level. pDCs displayed a distinct transcriptional profile, with upregulation of genes linked to metabolic and redox regulation, including *SLC25A5, GAPDH,* and *GSTP1*, together with immune-regulatory and cytoskeleton-associated genes such as *LGALS1, GRN, LSP1,* and *ARPC1B*. pDCs also showed downregulation of *PRKCH* and *ABR*, pointing to altered intracellular signaling and Rho GTPase-mediated cytoskeletal regulation. In NK cells, downregulated genes were concentrated in transcriptional regulators and homing-associated factors including *BACH2*, *TCF7*, and *CCR7*, consistent with a shift toward more terminally differentiated effector states (**Figure 4D**).

Pathway analysis revealed convergent themes across cell types (**Figure 4E**). pDCs and NK cells were enriched for mitochondrial respiration, mitochondrial translation, translation/elongation, and antigen presentation pathways, pointing to a state of heightened biosynthetic and metabolic activity in innate compartments. Conversely, Th17 and Th1/Th17 cells showed depletion of pathways involved in chromatin organization, SUMOylation, TGF-β/SMAD signaling, and cytokine signaling, indicating broad attenuation of transcriptional and regulatory programs in adaptive subsets in chronic patients (**Figure 4E**).

To explore whether transcriptional differences were reflected at the protein level, we assessed expression of TGFBR2, IκBα, and CD69 by flow cytometry across non-classical monocytes, NK cells, naïve CD4J T cells, and Th17 cells at six months post-infection. Although consistent directional trends were observed, notably higher IκBα expression in non-classical monocytes and naïve CD4J T cells, and reduced TGFBR2 in NK cells and non-classical monocytes in chronic patients, consistent with the transcriptomic depletion of TGF-β/SMAD signaling pathways, none reached statistical significance, likely reflecting limited statistical power (**Supplementary Figure 7**).

Together, these findings indicate that chronic patients at six months post-infection display persistent immune remodeling, characterized by metabolic and effector-associated pathway activity in innate compartments alongside reduced transcriptional and signaling pathway activity in selected adaptive T-cell subsets.

### Chronic CHIKV disease is associated with mistimed innate–adaptive communication networks

To assess whether transcriptional and compositional immune differences translated into altered intercellular communication, we performed ligand–receptor–based cell–cell interaction analysis using CellChat on scRNA sequencing data from acute infection and six months post-infection [34].

During acute infection, chronic and non-chronic patients exhibited distinct predicted ligand–receptor interaction networks. Acute non-chronic patients were characterized by myeloid-centered signaling, with classical monocytes acting as dominant hubs and increased outgoing communication toward CD8J T cell subsets and other myeloid populations, consistent with coordinated innate-to-adaptive immune priming. In contrast, acute chronic patients showed enhanced signaling from naïve CD4J T cells toward multiple CD8J T cell subsets, including naïve, central memory, and effector populations, indicating a shift away from myeloid-driven immune organization (**Figure 5A**).

**Figure 5:**
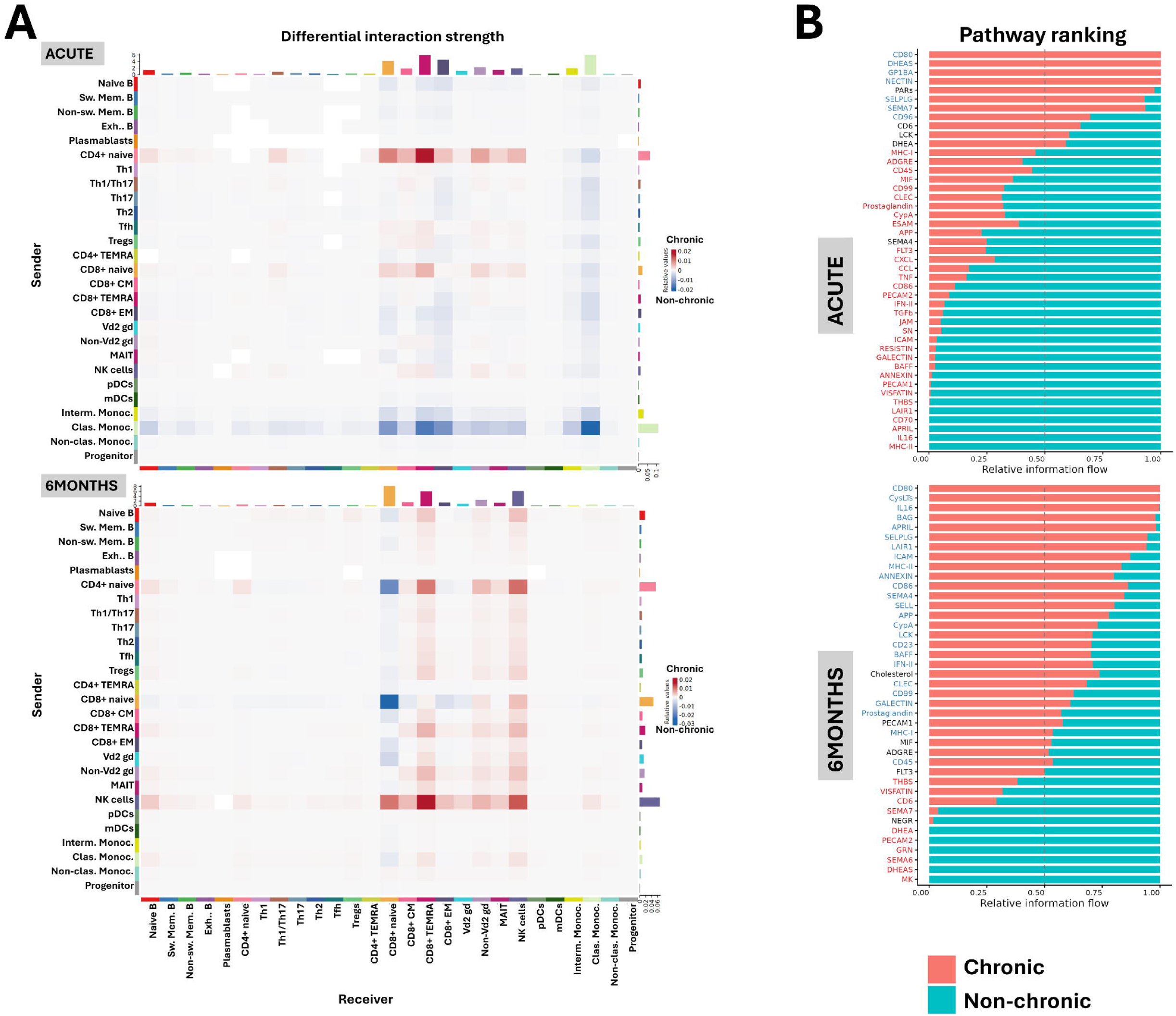
Distinct intercellular communication programs underlie chronic and non-chronic CHIKV trajectories. Ligand–receptor–based cell–cell communication analysis was performed using CellChat on scRNA sequencing data from acute infection and six months post-infection **(A)** Heatmaps showing differential interaction strength between sender (rows) and receiver (columns) cell populations. Red indicates higher communication probability in non-chronic patients, blue indicates higher communication probability in chronic patients. Only ligand–receptor interactions significant by CellChat permutation test (p ≤ 0.05) are included. **(B)** Pathway ranking plots displaying relative information flow per pathway and condition. Pathways are ordered by differences in relative information flow between groups. Statistical significance of pathway-level differences was assessed using CellChat rankNet (Wilcoxon signed-rank test with Benjamini–Hochberg correction), with FDR < 0.05 indicated by colored pathway labels.

At six months post-infection, these differences became more pronounced. Chronic patients displayed a prominent NK cell–centered communication pattern, marked by strong incoming signals to NK cells from multiple immune subsets, particularly naïve CD4J T cells and NK cells themselves, as well as increased outgoing signaling from NK cells toward naïve CD8J, CD8J TEM, CD8J T central memory (CM) and CD8J TEMRA cells and γδ T cells. This NK-centered pattern was absent in non-chronic patients, who instead showed stronger naïve CD4J and naïve CD8J T cell signaling toward naïve CD8J T cells, consistent with a more homeostatic adaptive network (**Figure 5A**).

Next, we evaluated differentially activated communication pathways across patient groups and timepoints. Ranking by relative information flow revealed a temporal inversion in 18 pathways, which were more active in non-chronic patients during acute infection but became relatively enriched in chronic patients at six months (**Figure 5B**). These included pathways involved in antigen presentation, T-cell co-stimulation, B-cell activation, adhesion/migration, and innate/inflammatory signaling, including MHC-I/MHC-II, CD86, APRIL/BAFF, ICAM/CD99, CLEC, IFN-II, and prostaglandin signaling.

Conversely, only a few pathways showed opposing or persistent patterns. SEMA7 and DHEAS signaling were higher in chronic patients during acute infection but not at six months, while VISFATIN, THBS, and PECAM signaling remained higher in non-chronic patients across both timepoints. SELPLG and CD80 were the only pathways persistently enriched in chronic patients. Notably, inflammatory and antiviral pathways, including TGF-β, TNF, CCL, and CXCL signaling, were selectively enriched in acute non-chronic patients, supporting stronger early immune activation in patients who recovered (**Figure 5B and Supplementary Figure 8**).

Cell-type–resolved pathway analysis further showed that these pathway-level differences reflected shifts in the cellular sources and targets of signaling rather than uniform pathway activation across all cell types. In acute non-chronic patients, classical monocytes contributed prominently to antigen-presentation, adhesion, and immune-signaling pathways, whereas these monocyte-associated interactions were reduced in chronic patients. At six months, chronic patients showed increased NK-cell and CD8J TEMRA involvement in inferred signaling networks, while non-chronic patients showed a more restricted adaptive pattern dominated by naïve CD8J T cells (**Supplementary Figure 8**).

Together, these results suggest that chronic CHIKV disease may be associated with reduced early myeloid-mediated immune priming during acute infection, resulting in altered T cell– and NK cell–centered predicted communication networks that persist alongside incomplete restoration of immune homeostasis.

### Longitudinal Transcriptomic Divergence Reveals Durable Immune Rewiring in Chronic CHIKV Disease

To characterize disease-specific recovery dynamics, we quantified differences in longitudinal gene expression changes between chronic and non-chronic patients using ΔlogJFC values across cell types ([C_A_ – C_6M_] − [NC_A_ – NC_6M_]). ΔlogJFC therefore measures how strongly gene expression trajectories differ between groups from the acute to the six months timepoint. Genes with ΔlogJFC values near zero follow similar recovery trajectories, whereas larger absolute values indicate divergent temporal regulation. Pathway enrichment was performed on ΔlogJFC genes to identify biological programs associated with these trajectory differences. To prioritize affected cell types, we ranked them by the number of genes with |ΔlogJFC| ≥ 1 and performed pathway enrichment on the top-ranked cell subsets (**Figure 6A).**

**Figure 6.**
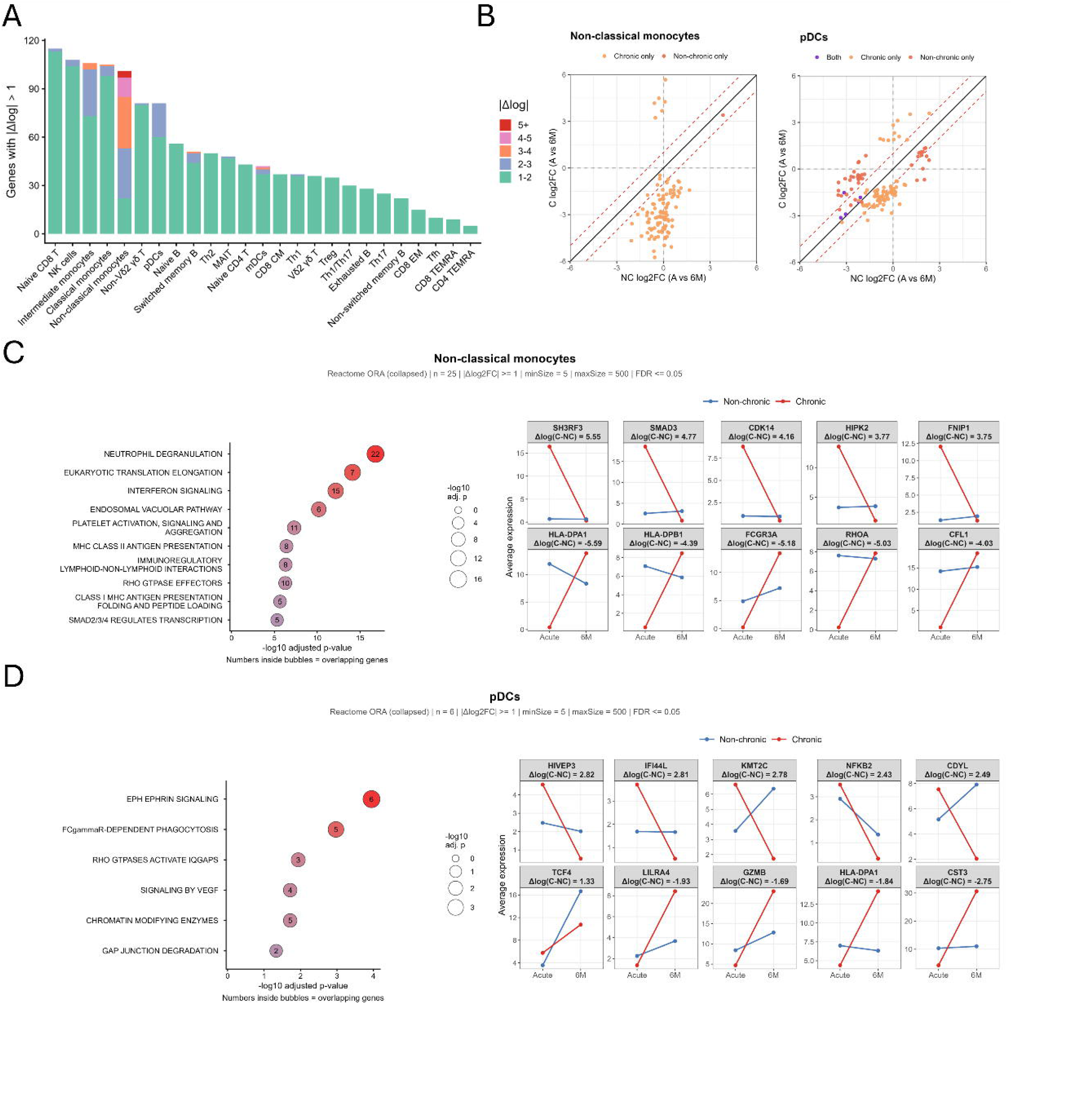
Longitudinal Transcriptomic Divergence Highlights Maladaptive Recovery Trajectories in Chronic CHIKV Disease. **(A)** Stacked bar chart showing the number of DEGs with |Δlog□FC| ≥ 1 per immune cell type, ranked from highest to lowest. Bars are colored by Δlog□FC magnitude bins (1–2, 2–3, 3–4, 4–5, ≥5). **(B) S**catter plots comparing longitudinal log□FC values (Acute - six months) between chronic (C, y-axis) and non-chronic (NC, x-axis) patients for non-classical monocytes (left) and pDCs (right). Each dot represents a gene significantly differentially expressed in at least one of the two longitudinal comparisons. Colors denote genes with significant trajectory divergence in chronic patients only (orange), non-chronic patients only (blue) or both groups (purple). Dashed red lines indicate |log□FC| = 1 thresholds. Genes near the diagonal indicate similar recovery dynamics between groups, whereas deviation from the diagonal reflects differences in longitudinal expression trajectories between chronic and non-chronic patients. **(C-D)** Pathway enrichment analysis was performed using Reactome overrepresentation analysis (ORA, collapsed) for non-classical monocytes (C) and pDCs (D). Significantly enriched pathways are shown in the left panels and ranked by −log10 adjusted p-value. Bubble size reflects −log10 adjusted p-value, and numbers inside bubbles indicate the number of overlapping genes. Selected genes with high Δlog□FC values are shown as trajectory line plots (right), with non-chronic patients in blue and chronic patients in red. Δlog values are indicated per gene panel. Pathway enrichment was performed using Reactome ORA on genes with |Δlog□FC| ≥ 1 against a background of all genes tested in the scRNA sequencing differential expression analysis (minSize = 5, maxSize = 500, FDR ≤ 0.05). Redundant pathways were removed by iteratively discarding any pathway with a Jaccard similarity ≥ 0.60 or smaller-set overlap ≥ 0.80 with a higher-ranked pathway, applied to the top 40 significant terms per cell type. Pathogen-specific Reactome terms unrelated to CHIKV were excluded prior to analysis.

Non-classical monocytes, intermediate monocytes and pDCs harbored the greatest number of genes with |ΔlogJFC| ≥ 2, identifying innate immune compartments as the primary contributors of divergent recovery trajectories between chronic and non-chronic patients (**Figure 6A–B**). Beyond these myeloid populations, conserved trajectory divergence was also detected across multiple cytotoxic and effector T cell subsets, including mucosal-associated invariant T (MAIT) cells, γδ T cells and CD8J T cell populations, indicating that immune remodeling in chronic disease extends across both innate and adaptive compartments.

Pathway enrichment analysis in non-classical monocytes revealed prominent divergence in interferon signaling, Rho GTPase signaling, antigen processing and presentation, and neutrophil degranulation pathways (**Figure 6C**). To visualize the direction of the underlying longitudinal differences, we plotted representative genes with strong ΔlogJFC values and/or biological relevance to these pathways. Genes with the strongest trajectory divergence included *HLA-DPA1* and *HLA-DPB1* (antigen presentation), *FCGR3A/CD16* (Fc receptor-mediated effector function), *RHOA* and *CFL1* (cytoskeletal regulation), *CDK14* (Wnt-associated signaling), *FNIP1* (metabolic signaling), *SH3RF3* and *HIPK2* (intracellular signaling scaffolds), and *SMAD3*, the central transcriptional effector of TGF-β signaling (**Figure 6C**). Notably, *HLA-DPA1, HLA-DPB1, FCGR3A*, *RHOA* and *CFL1* showed increasing expression toward six months in chronic patients, whereas expression remained stable or declined over time in non-chronic individuals, highlighting persistent remodeling of antigen presentation, Fc receptor signaling and cytoskeletal regulation.

Similarly, pDCs displayed divergent longitudinal regulation of pathways linked to Rho GTPase signaling, Eph/ephrin-mediated signaling, Fcγ receptor-dependent phagocytosis and VEGF signaling (**Figure 6D**). Representative trajectory genes again showed outcome-specific temporal regulation. Genes with the highest ΔlogJFC values included *IFI44L* (interferon-stimulated antiviral response), *NFKB2* (NF-κB inflammatory signaling), *GZMB* (cytotoxic effector granzyme B), *HIVEP3*, *KMT2C*, and *CDYL* (transcriptional and chromatin regulation), *HLA-DPA1* (antigen presentation) and *CST3* (lysosomal/protease regulation), alongside *TCF4*, a key transcription factor for pDC lineage identity and survival, and *LILRA4*, an inhibitory receptor characteristic of pDCs (**Figure 6D)**. *IFI44L* and *NFKB2* declined more strongly over time in chronic patients, suggesting altered resolution of antiviral and inflammatory signaling. In contrast*, LILRA4, GZMB, CST3* and *HLA-DPA1* increased toward six months in chronic patients, whereas *TCF4* increased in both groups but more prominently in non-chronic patients. *KMT2C* and *CDYL* showed opposite trajectories over time between chronic and non-chronic patients, decreasing over time in the former. This pattern suggests that pDC-associated recovery trajectories diverge between outcomes, with non-chronic patients showing stronger restoration of canonical pDC identity programs, whereas chronic patients show greater remodeling toward inhibitory, cytotoxic and antigen-presentation-associated states (**Figure 6D**).

Intermediate monocytes similarly exhibited altered longitudinal regulation of interferon signaling, PD1 signaling, translational pathways and chaperone-mediated autophagy. As observed in non-classical monocytes, many genes followed divergent temporal trajectories between chronic and non-chronic patients, further emphasizing widespread disruption of coordinated innate immune resolution (**Supplementary Figure 9**).

Across innate immune populations, recurrent enrichment of MHC class II-associated pathways, Fc receptor signaling and phagocytosis-related pathways identified antigen presentation remodeling as a central feature of chronic disease evolution. In parallel, enrichment of Rho GTPase and Eph/ephrin signaling pathways suggested broader alterations in cytoskeletal organization, immune cell interaction and migratory programs during chronic disease progression.

To identify conserved molecular features associated with chronic disease, we next examined genes displaying trajectory divergence across multiple immune cell types. Genes present in at least six cell types with |ΔlogJFC| ≥ 1 included *ACTG1, HLA-DPA1, HSPA8, HOPX, S100A4, RICTOR, TRIP12, MECP2, BIRC3* and *IFI44L* (**Supplementary Figure 10**). These genes showed remarkably consistent temporal behavior across immune compartments (**Supplementary Figure 11**), suggesting coordinated systemic immune remodeling rather than isolated cell type-specific dysregulation. Functionally, these conserved genes represented multiple biological programs, including antigen presentation (*HLA-DPA*1), interferon signaling (*IFI44L*), cytoskeletal organization and migration (*ACTG1, S100A4*), inflammatory and survival signaling (*BIRC3*), and mTORC2-dependent metabolic regulation (*RICTOR*).

Together, these findings support a model in which chronic CHIKV disease is characterized not by persistent broad inflammatory activation, but by coordinated and durable rewiring of immune recovery trajectories across multiple immune compartments, particularly within innate myeloid populations.

## Discussion

Our longitudinal immune profiling analysis suggests that chronic CHIKV disease is not driven by persistent systemic inflammation, but is associated with coordinated failures in immune network organization that begin during acute infection and evolve into durable transcriptional, metabolic, and cellular reprogramming that fails to resolve by six months post-infection (**Figure 7**). Patients who progressed to chronic disease exhibited early immune hypoactivation, characterized by reduced interferon responses, dampened inflammatory cytokine production, and impaired activation of both innate and adaptive immune compartments, followed by sustained remodeling of innate immune populations, including non-classical monocytes, pDCs and NK cells, that persisted into convalescence.

**Figure 7:**
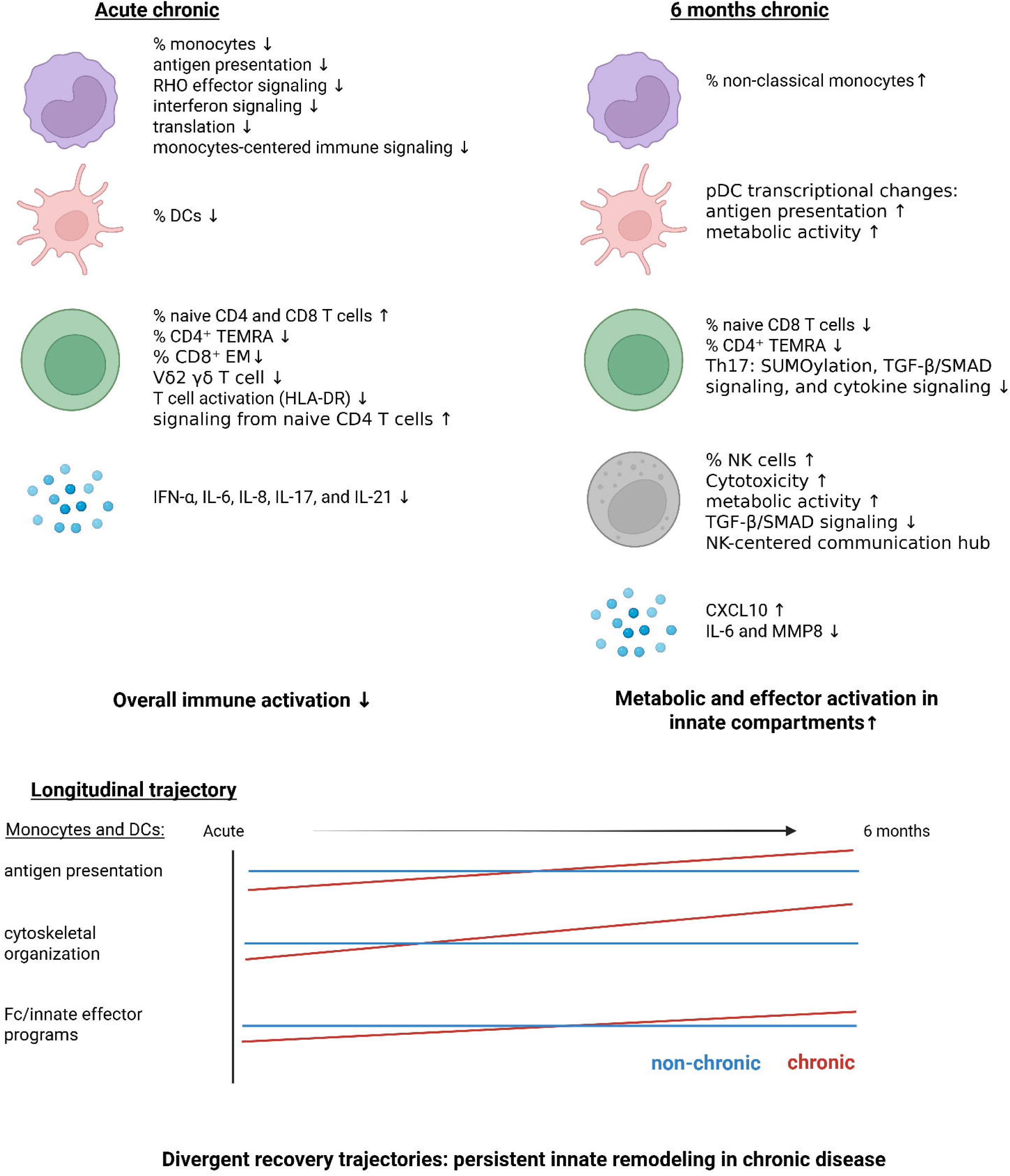
Overview of immune alterations associated with chronic chikungunya disease. Chronic patients showed dampened acute immune activation, with reduced innate cell abundance/function, lower T cell activation, and decreased inflammatory cytokines. At six months, chronic disease was characterized by persistent innate remodeling, including increased non-classical monocytes, pDCs and NK cell metabolic/effector changes, elevated CXCL10, and reduced IL-6/MMP8. Longitudinally, chronic patients displayed sustained divergence in antigen presentation, cytoskeletal organization, and Fc/innate effector programs compared with non-chronic patients.

During acute infection, chronic patients showed attenuation of innate and adaptive immune activation, including reduced systemic frequencies of classical, intermediate, and non-classical monocytes as well as mDCs and pDCs, significantly lower IFN-α, IL-6, IL-8, IL-17 and IL-21, diminished monocyte transcriptional activation, increased naïve T cell populations, and decreased antigen presentation.

Previous studies have described complementary immune evasion mechanisms employed by CHIKV that further support this hypoactivation: CHIKV inhibits NF-κB and Wnt signaling through nsP2 and nsP3 [36–38], reduces surface MHC-I expression [39], and downregulates antiviral genes including *LIFR* and *MMP8*, as well as neutrophil degranulation and MHC-I antigen processing pathways [39, 40]. The reduction in LIFR may be mechanistically relevant because LIFR can promote IL-6 production through STAT signaling [41]. Its downregulation may therefore contribute to reduced IL-6 production and broader attenuation of early cytokine-mediated immune activation. Because IFN-α is critical for limiting early viral replication, and IL-6 contributes to early antiviral immune coordination [42], their combined attenuation could potentially contribute to incomplete viral clearance and extend the window during which CHIKV antigens interface with the host immune system. Although data on CHIKV persistence remain inconclusive, with one study reporting CHIKV RNA/protein in synovial macrophages up to 18 months post-infection [10] and another detecting no viral RNA in synovial fluid [15], a recent mouse model demonstrated that ongoing viral replication in joint-associated macrophages drives inflammatory macrophage and CD4J T cell accumulation, and that inhibiting replication reduced joint inflammation [14]. These findings do not exclude transient or tissue-restricted antigen persistence as a contributor to later chronic symptoms. Together, these acute features align with Colombian cohort data showing that low early cytokine levels predict chronic outcomes [43], and support the emerging concept that early immune hypoactivation, rather than excessive inflammation, is a key determinant of chronicity.

At the network level, classical monocytes failed to establish themselves as communication hubs, with key antigen presentation and immune-licensing pathways largely absent, including MHC-II, APRIL, IL-16, and adhesion-related signaling. In contrast, non-chronic patients displayed robust classical monocyte-centered signaling toward CD8J T cells and myeloid populations, enriched for antigen presentation, adhesion, and inflammatory pathways, consistent with effective innate-to-adaptive immune priming. Notably, chronic patients exhibited a shift toward naïve CD4J T cell–centered communication toward multiple CD8J subsets in the absence of robust myeloid licensing, suggesting altered rather than fully effective adaptive activation. These findings align with studies showing that a strong early innate response facilitates viral clearance [44–46], whereas low early cytokine levels predict chronic outcomes [43].

By six months post-infection, most systemic adaptive cytokine differences had resolved, whereas innate immune alterations persisted. Chronic patients showed higher frequencies of non-classical monocytes, elevated CXCL10, and transcriptional remodeling in pDCs and NK cells. These three populations emerged as a central innate axis of chronic disease. Non-classical monocytes and pDCs showed the strongest longitudinal transcriptional divergence, while NK cells dominated the predicted communication network at six months. These findings could be compatible with the broader framework of innate immune memory, in which innate cells can acquire durable functional states ranging from trained hyperresponsiveness to tolerance-like hyporesponsiveness through related epigenetic and metabolic mechanisms [47].

The acute profile of chronic progressors resembled a tolerance-like innate state, with reduced NF-κB, interferon, antigen-presentation and inflammatory signaling. However, this state did not persist as uniform immune suppression. Instead, chronic patients later displayed renewed antigen-presentation, metabolic and cytotoxic programs, together with NK-centered predicted communication. Altered NF-κB-related signaling, increased IκBα trends by flow cytometry, and divergent *SMAD3* and *HIPK2* regulation in non-classical monocytes further support a model in which early regulatory or tolerance-like programming evolves into persistent innate immune remodeling. Both genes were already more highly expressed in chronic patients during acute infection and followed divergent longitudinal trajectories. Because *HIPK2* can potentiate TGF-β/SMAD3 transcriptional activity, and TGF-β/SMAD signaling can restrain NF-κB-driven inflammation through mechanisms including IκBα induction [48, 49], these findings suggest an early bias toward TGF-β/SMAD-associated regulatory programming in chronic progressors.

Non-classical monocytes and pDCs showed the most extensive longitudinal divergence, with sustained ΔlogJFC shifts and broad transcriptional rewiring of interferon signaling, antigen presentation, Fc effector programs and cytoskeletal organization. In parallel, *S100A4* upregulation in non-classical monocytes of chronic patients over time, points to a fibro-inflammatory component of the chronic innate signature, linking persistent monocyte remodeling to tissue-remodeling pathways implicated in synovial inflammation and chronic arthralgia [50]. Reports of expanded intermediate and non-classical monocyte subsets in acute and chronic CHIKV [51] further support non-classical monocytes involvement in chronic pathogenesis. The transcriptional profile of pDCs at six months is consistent with a persistently activated yet functionally remodeled state characterized by inhibitory signaling and enhanced antigen-presentation programs. NK cell remodeling was characterized by downregulation of *BACH2*, *CCR7* and *TCF7* together with increased cytotoxic effector signatures. Because *BACH2* restrains NK cell maturation and effector function, *CCR7* regulates lymphoid homing, and *TCF7* supporting survival during maturation, this pattern suggests terminal effector skewing with loss of homing- and persistence-associated programs [52–54]. The dominance of NK cells in the convalescent communication network of chronic patients may therefore reflect sustained cytotoxic innate activity rather than productive immune resolution.

Across multiple T cell subsets, *RICTOR* showed pronounced ΔlogJFC decline in chronic patients. Given mTORC2’s essential role in metabolism, cytoskeletal reorganization, and effector-to-memory transitions [55, 56], its attenuation provides a mechanistic basis for impaired T cell recovery, a pattern shared with mTOR dysregulation in T cell exhaustion during chronic viral infections such as HIV and HCV [56].

The persistence of elevated CXCL10 at six months suggests incomplete resolution of interferon-associated inflammation, despite the absence of measurable circulating IFN-α in plasma. CXCL10 is an interferon-inducible chemokine responsive to type I and type II interferon pathways [57]. Elevated CXCL10 may therefore reflect residual interferon-stimulated activity driven by low-level, tissue-localized, or IFN-γ–associated signaling rather than a detectable systemic IFN-α response. Potential sources of such signaling include activated NK or T cells, local tissue inflammation, or ongoing antigenic stimulation. This interpretation is further supported by the disproportionate longitudinal decline of *IFI44L*, a negative regulator of interferon signaling [58], across multiple cell types in chronic patients, suggesting impaired re-establishment of interferon negative-feedback loops and persistence of partially activated antiviral programs. Because CXCL10 normally decreases after the acute viremic/seroconversion phase of CHIKV infection [16, 59], its persistence at six months may mark incomplete resolution of interferon-associated inflammation. The emergence of NK-centered communication networks is consistent with this model, as CXCL9/10/11–CXCR3 chemokine axes direct the migration of activated T cells and NK cells [57, 60], creating a potential feed-forward loop of sustained innate immune recruitment and incomplete resolution.

A key insight from the intercellular communication analysis is that chronic CHIKV disease may involve mistimed immune coordination rather than sustained inflammation alone. Pathways linked to antigen presentation, B-cell support, leukocyte trafficking and T cell priming were prominent during acute infection in non-chronic patients, but became relatively more active in chronic patients only at six months, suggesting that antigen-processing and immune-licensing programs are not absent in chronic patients but temporally displaced. Such late reactivation may reflect persistent antigenic stimulation, unresolved tissue inflammation or compensatory activation after inadequate early priming. This raises the outstanding question of which antigens are being presented at six months, persisting CHIKV-derived antigens, cross-reactive self-antigens, or bystander antigens released during tissue inflammation, which will require synovial profiling, antigen-specific T cell assays or immunopeptidomics to resolve. Earlier studies have highlighted chronic IL-6, IL-8, and IL-17 signatures [18, 19, 59], but we did not detect such cytokines in plasma, flow cytometry, or scRNA sequencing at six months. A Colombian cohort at 22 months [15, 43] similarly found no persistent inflammatory cytokines in plasma or viral RNA in synovial fluid, indicating that systemic inflammation is not universally associated with chronic CHIKV.

Although our analysis provides detailed insight into systemic immune trajectories, several limitations remain. We did not perform BCR sequencing or comprehensive autoantibody profiling, leaving open the possibility of autoreactive responses in a subset of patients. Systemic responses measured in PBMCs may not fully represent immune dynamics within joint tissues, the primary site of pathology. The flow cytometry cohort was modest, limiting detailed assessment of rare populations and statistical power. Finally, we were unable to quantify viral RNAemia during the acute phase of infection or determine the timing of sampling at the acute phase of infection, as inclusions were in response to the CHIKV epidemic. Even though RNAemia is short (between -2 and 8 days after onset of fever) [61], discordant sampling could affect our findings.

The early network-level dysfunction identified here suggests a critical intervention window during acute infection, before chronic disease is established. Strategies enhancing monocyte antigen presentation capacity or restoring innate-adaptive coordination during the first week of infection could plausibly reduce chronic disease development. In established chronic disease, the pathways implicated here, such as interferon feedback mechanisms, the HIPK2/SMAD3/TGF-β/NF-κB circuit, mTORC2-mediated metabolic remodeling and FcγR signaling, suggest therapeutic strategies aimed at restoring immune homeostasis rather than broadly suppressing inflammation.

## Declaration of Interest

The authors declare no competing interests.

## Acknowledgements

We would like to thank Dr. Rithea Leang and Dr. Huy Rekol from the National Center for Parasitology, Entomology and Malaria Control for their support. Biorender was used to design figures.

## Funding

This work was supported by Wellcome Trust grant no. 311543/Z/24/Z. The Cantaert lab is additionally supported by the NIH (PICREID,1U01AI151758, R01 AI175134, and R01 AI137276) and Horizon Europe (grant no. 101137033). LDC was supported by a post-doctoral grant Calmette & Yersin of the Pasteur Network.

## Author Contribution

Conceptualization: TC, LDC

Methodology: LDC, GG, TC

Investigation: LDC, CB, SLay, BH, SK

Formal Analysis: LDC, GG

Resources (Sample collection): SS, Sly

Data Curation: LDC, AM

Writing – Original Draft: LDC, TC

Writing – Review & Editing: LDC, GG, CB, TC

Visualization: LDC, GG

Supervision: SLy, VD, TC

Project Administration: TC Funding Acquisition: TC

## Data and code availability

Any additional information required to reanalyze the data reported in this paper is available from the lead contact upon request. Data will be deposited upon acceptance of the manuscript.

## Supplemental information

Document S1: Figure S1-S11, Table S1

## Notes

### Competing Interest Statement

The authors have declared no competing interest.

