## Supplemental data for "Early Immune Hypoactivation and Persistent Innate Reprogramming Characterize Chronic Chikungunya Disease"

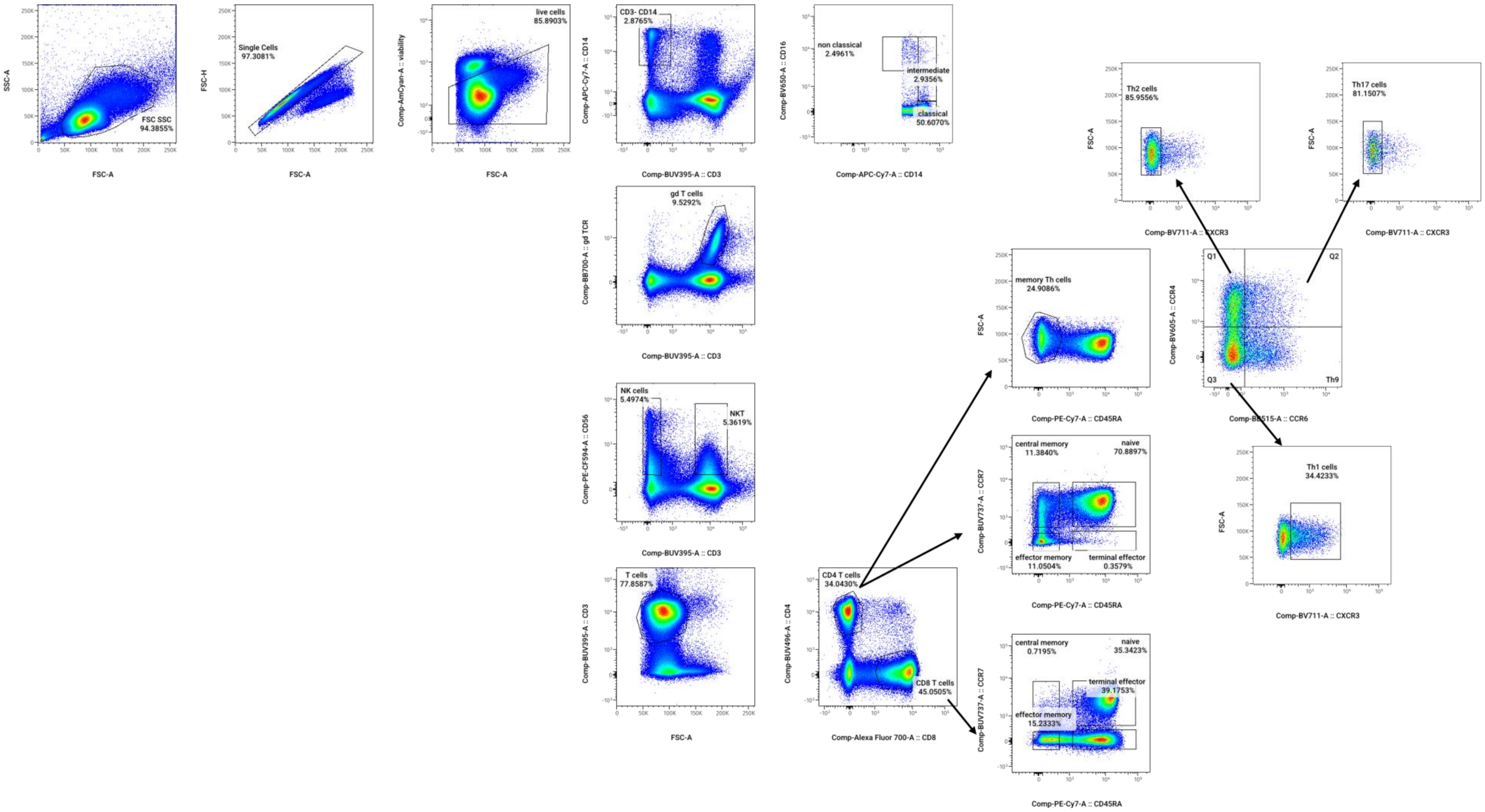


**Supplementary Figure 1: Flow cytometry gating strategy for innate and adaptive immune subsets.**

Cells were first gated on FSC–SSC to remove debris, followed by selection of singlets (FSC-H vs FSC-W) and viable cells using a live/dead discriminator. From total viable cells, monocytes were identified as CD3⁻/CD14⁺ and further subdivided into classical (CD3⁻/CD14⁺/CD16⁻), intermediate (CD3⁻/CD14⁺/CD16⁺), and non-classical (CD3⁻/CD14^dim/CD16⁺) subsets. γδ T cells were gated as CD3⁺/γδ⁺, while NK and NKT cells were defined as CD3⁻/CD56⁺ and CD3⁺/CD56⁺, respectively. T cells were gated as $CD3^+$ and subsequently separated into $CD4^+$ and $CD8^+$ populations. Within the $CD3^+CD4^+$ compartment, memory Th cells were defined as $CD45RA^-$, and Th subsets were discriminated based on chemokine receptor expression: Th1 ($CD45RA^-/CCR4^-/CCR6^-/CXCR3^+$), Th2 ($CD45RA^-/CCR4^+/CCR6^-/CXCR3^-$), Th9 ($CD45RA^-/CCR4^-/CCR6^+$), and Th17 ($CD45RA^-/CCR4^+/CCR6^+/CXCR3^-$). $CD4^+$ and $CD8^+$ T-cell memory subsets were further classified using CD45RA and CCR7 expression to identify naïve ($CD45RA^+/CCR7^+$), central memory ($CD45RA^-/CCR7^+$), effector memory ($CD45RA^-/CCR7^-$), and terminal effector ($CD45RA^+/CCR7^-$) populations. Representative plots illustrate each gating step used to derive frequencies for downstream analyses.

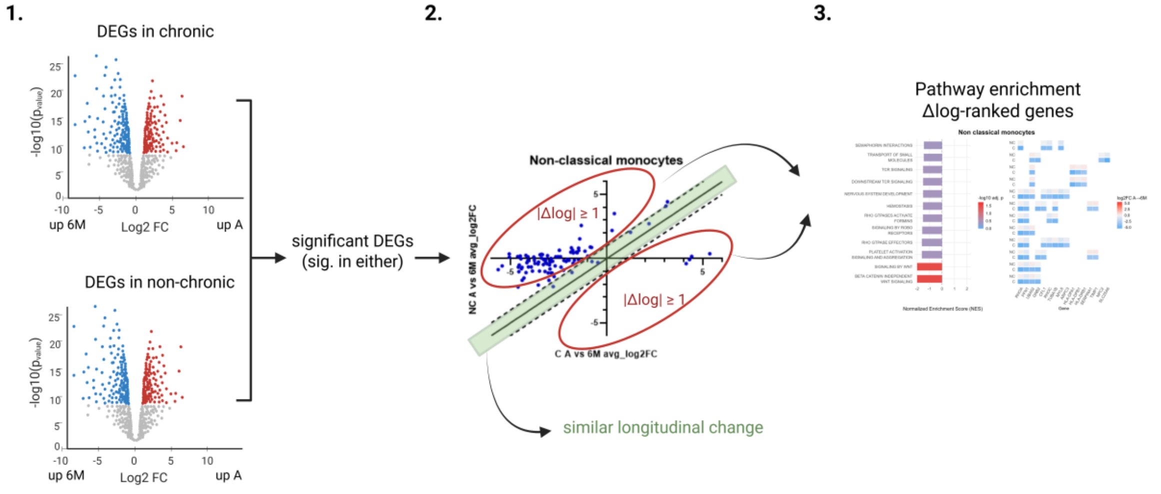


**Supplementary Figure 2. Workflow for identifying genes with divergent longitudinal expression trajectories.**

(1) Longitudinal differential expression analysis was performed separately in chronic and non-chronic patients (acute vs. 6 months). Genes significantly differentially expressed in at least one comparison were retained.

(2) Longitudinal expression changes were calculated as $\log_2$ fold change (acute → 6 months) for each group and plotted against each other to visualize recovery trajectories. Divergence between groups was quantified as

$\Delta\log_2FC = \log_2FC\_A{\rightarrow}6M(Chronic) - \log_2FC\_A{\rightarrow}6M(Non\text{-}chronic)$.

Genes with $|\Delta\log_2FC| \geq 1$ were considered to show substantial trajectory divergence.

(3) by $\Delta\log_2FC$ genes were used for Reactome pathway enrichment analysis (ORA) to identify biological programs associated with divergent recovery dynamics.

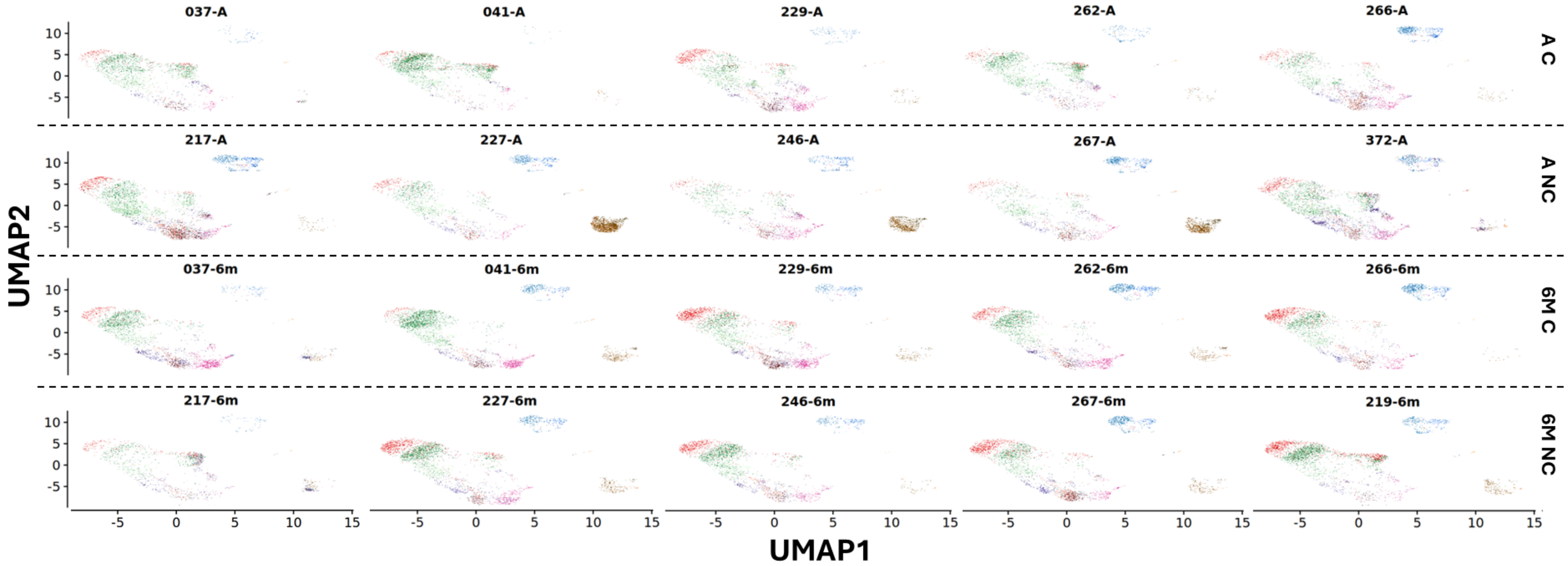


**Supplementary Figure 3: Per-sample UMAP and cell type composition of PBMC single-cell data.**
(A) UMAP representation of integrated PBMC single-cell RNA-seq data, shown separately for each sample and grouped by clinical outcome and timepoint: acute chronic (AC), acute non-chronic (ANC), 6 months chronic (6M C), and 6 months non-chronic (6M NC). Cells are colored by annotated cell type and projected onto a shared embedding following Harmony integration.

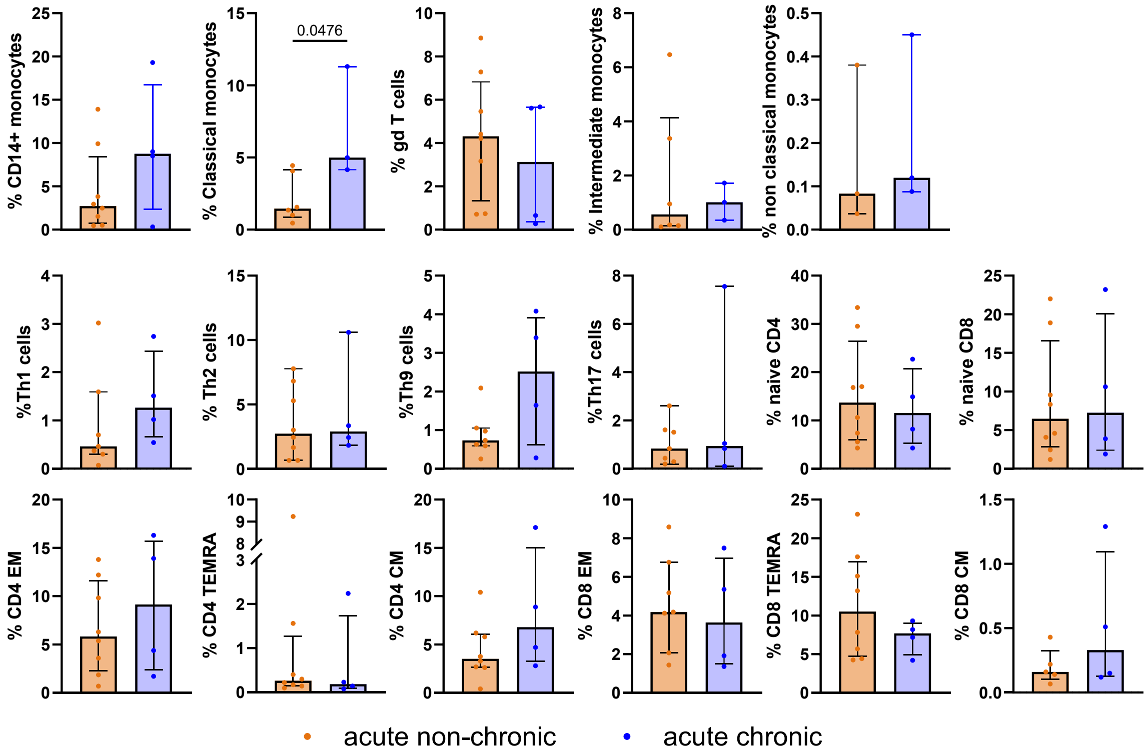


**Supplementary Figure 4: Flow cytometric analysis of acute-phase immune cell subset frequencies in chronic and non-chronic patients**

Flow-cytometry–derived frequencies of selected immune cell subsets among total viable cells during acute CHIKV infection, stratified by later clinical outcome as non-chronic or chronic disease. Monocytes were defined as $CD3^{-}CD14^{+}$ cells and further classified as classical monocytes ($CD3^{-}CD14^{+}CD16^{-}$), intermediate monocytes ($CD3^{-}CD14^{+}CD16^{+}$), and non-classical monocytes ($CD3^{-}CD14$^dim^$CD16^{+}$). Memory Th subsets were defined as memory Th1 ($CD3^{+}CD4^{+}CD45RA^{-}CCR4^{-}CCR6^{-}CXCR3^{+}$), Th2 ($CD3^{+}CD4^{+}CD45RA^{-}CCR4^{+}CCR6^{-}CXCR3^{-}$), Th9 ($CD3^{+}CD4^{+}CD45RA^{-}CCR4^{-}CCR6^{+}$), and Th17 cells ($CD3^{+}CD4^{+}CD45RA^{-}CCR4^{+}CCR6^{+}CXCR3^{-}$). $CD4^{+}$ and $CD8^{+}$ T-cell memory subsets were defined as effector memory T cells (TEM; $CD45RA^{-}CCR7^{-}$), terminally differentiated effector memory T cells re-expressing CD45RA (TEMRA; $CD45RA^{+}CCR7^{-}$), and central memory T cells (TCM; $CD45RA^{-}CCR7^{+}$), gated on $CD3^{+}CD4^{+}$ or $CD3^{+}CD8^{+}$ T cells, respectively. Each point represents an individual donor; bars indicate the median and error bars show the interquartile range. Comparisons between chronic and non-chronic patients were performed using the Mann–Whitney U test. Sample sizes ranged from n = 3–8, depending on the subset analyzed.

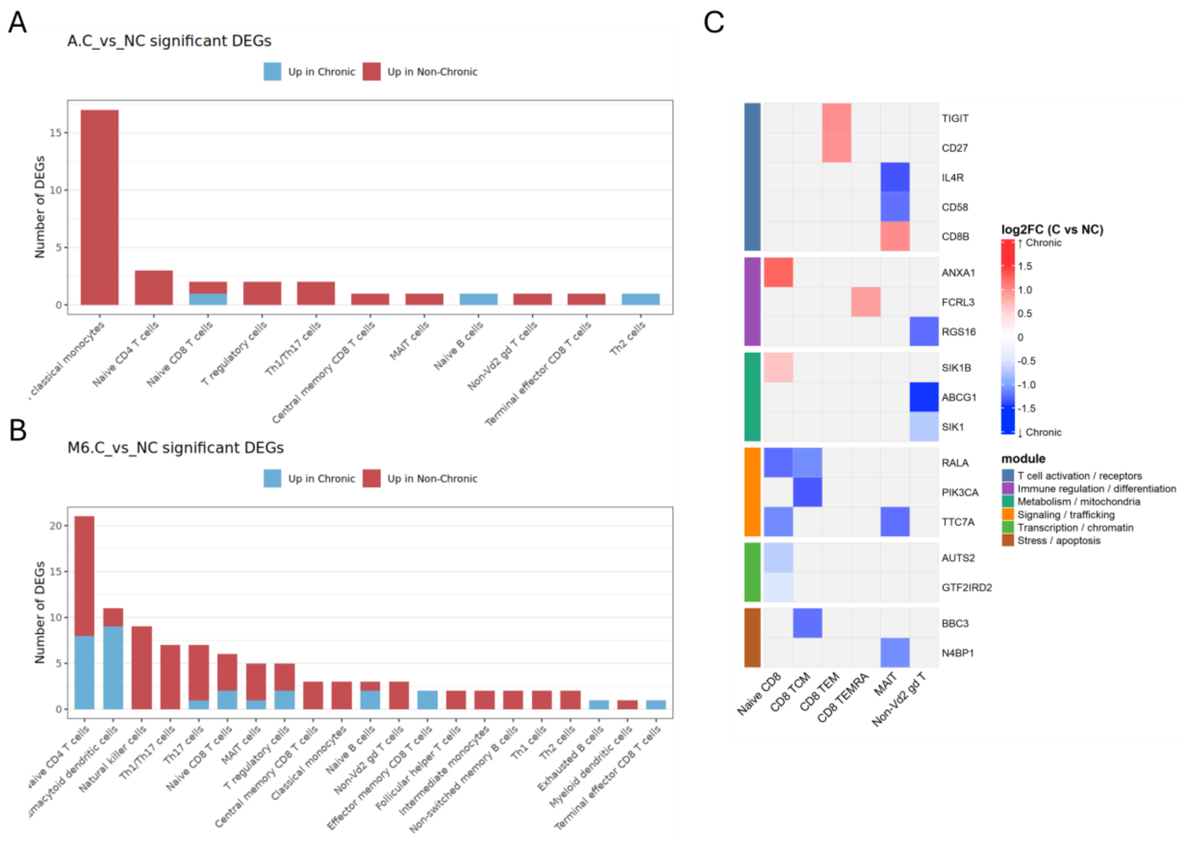


**Supplementary Figure 5: Distribution of significant differentially expressed genes across immune cell subsets.**

**(A-B)** Stacked bar plots showing the number of significant differentially expressed genes (DEGs) detected in each annotated immune cell subset during the acute phase (A.C_vs_NC)**(A)** and at 6-month follow-up (M6.C_vs_NC)**(B)**. Bars indicate the total number of significant DEGs per cell type, stratified by direction of regulation: genes upregulated in chronic patients are shown in blue, and genes upregulated in non-chronic patients are shown in red.

**(C)** Heatmap of DEGs across CD8 T cell immune subsets, MAIT and non-Vδ2 γδ T cells at 6 months. DEGs were defined as |log2FC| > 0.25 (chronic vs non-chronic) and grouped into co-regulated functional modules. Color scale indicates log2 fold change, with red representing higher expression in chronic patients and blue representing lower expression.

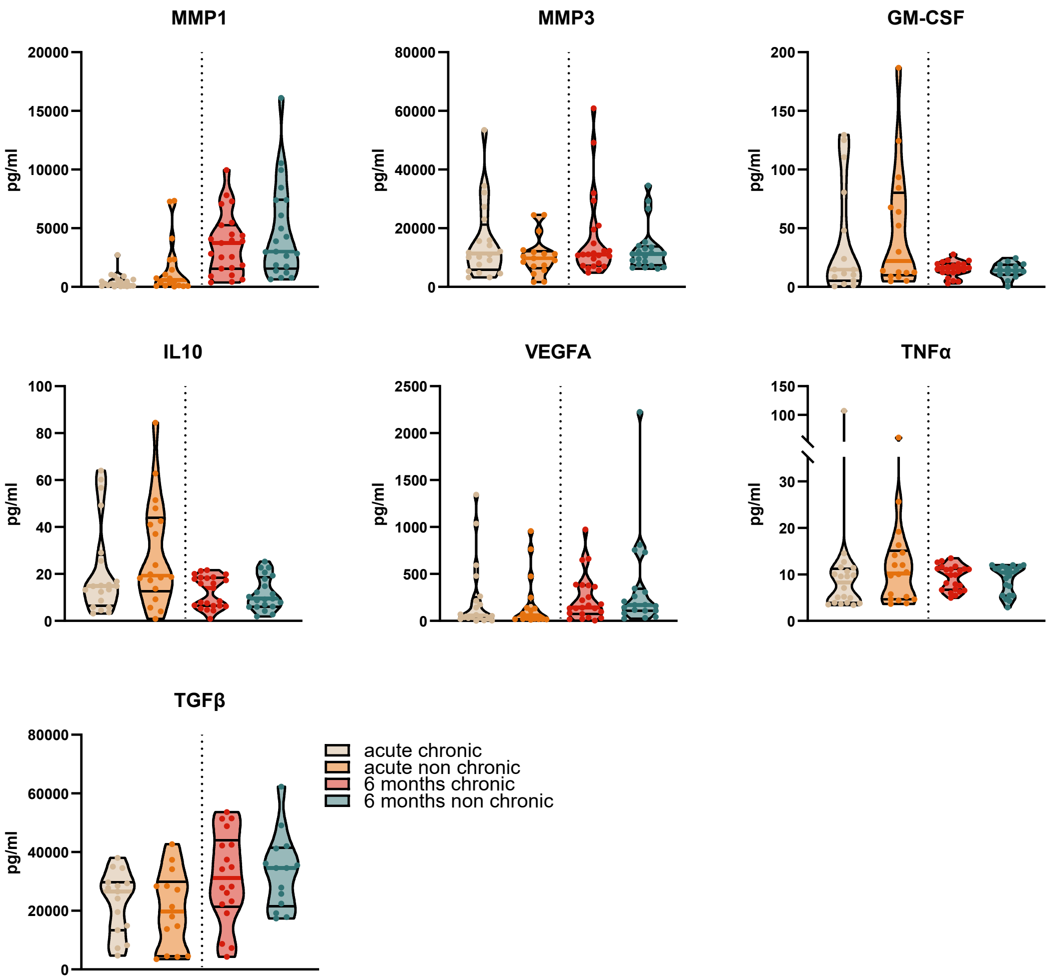


**Supplementary Figure 6. Additional cytokines and interleukins measured.**
Violin plots display concentrations of additional cytokines and interleukins (MMP1, MMP3, GM-CSF, IL10, VEGFA, TNFα, and TGFβ) measured during the acute and 6-month timepoints in chronic and non-chronic CHIKV patients. Individual donor values are shown for each group. Group comparisons were performed using the Mann–Whitney U test.

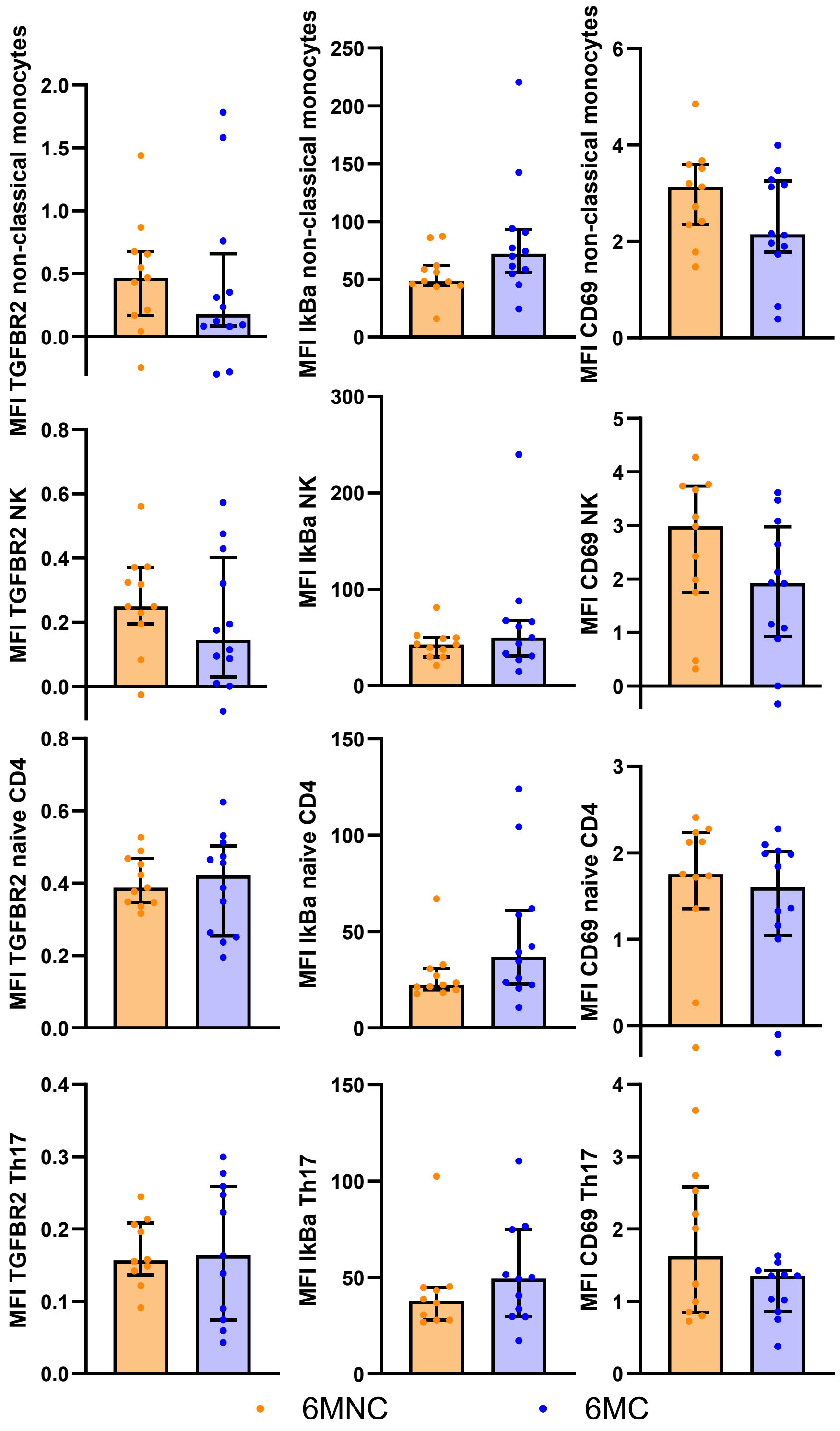


**Supplementary Figure 7: Flow cytometric assessment of selected inflammatory and regulatory markers at 6-month follow-up.**

Bar plots showing normalized median fluorescence intensity (MFI) of TGFBR2, IκBα, and CD69 in non-classical monocytes, NK cells, naïve CD4 T cells, and Th17 cells at 6 months after acute CHIKV infection. Patients are stratified according to clinical outcome as non-chronic (orange) and chronic (blue). Each dot represents an individual patient; bars indicate the median and error bars show the interquartile range. P-values were calculated using a two-sided Mann–Whitney U test and are indicated above each comparison.

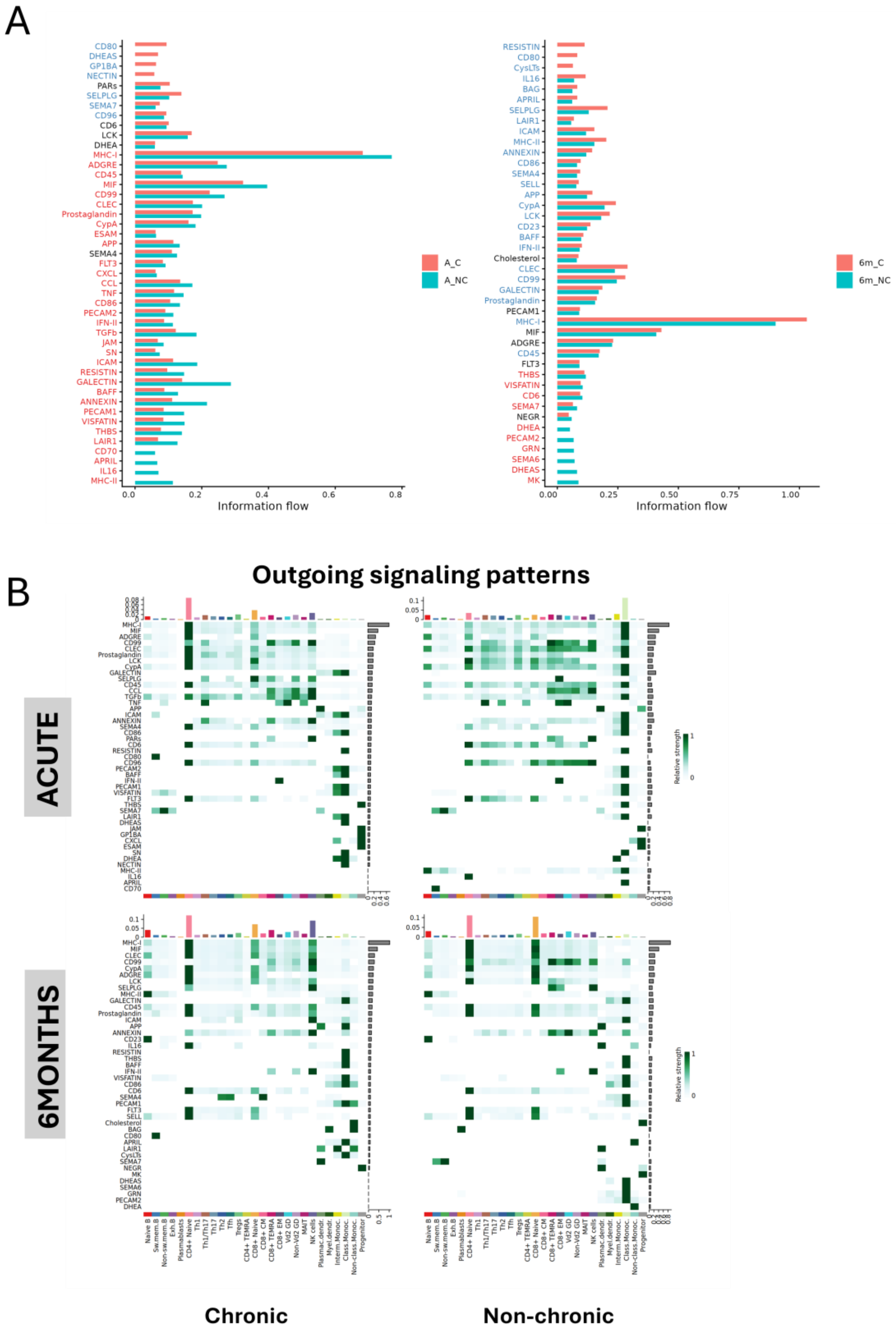


**Supplementary figure 8: CellChat analysis of inferred communication pathway activity across disease outcomes and timepoints.**

**(A)** Total information flow of inferred ligand–receptor signaling pathways in acute chronic (A_C), acute non-chronic (A_NC), six-month chronic (6m_C), and six-month non-chronic (6m_NC) patients. Bar plots compare pathway-level information flow between chronic and non-chronic patients during acute infection and at six months post-infection.

**(B)** Pathway-by-cell-type heatmaps showing sender-cell contributions to inferred outgoing signaling activity for each condition. Rows represent communication pathways and columns represent sender cell populations. Color intensity indicates the relative contribution of each sender cell type to the indicated pathway within each condition.

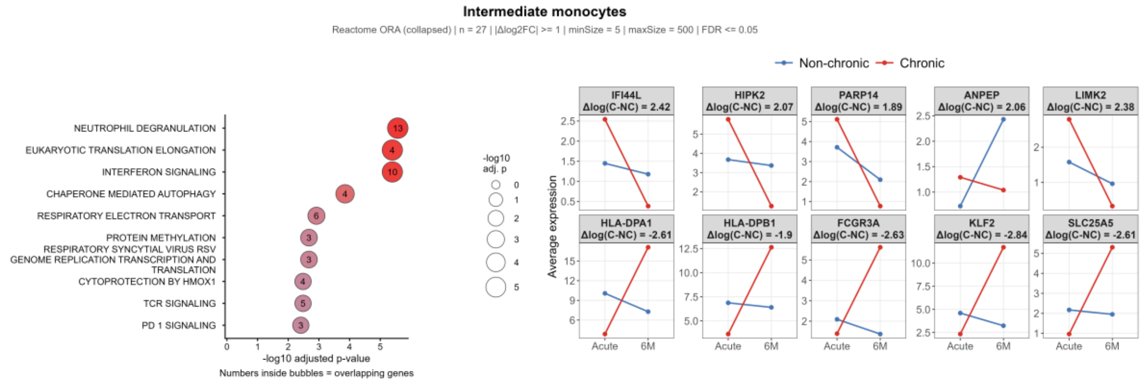


**Supplementary Figure 9: Longitudinal pathway divergence in intermediate monocytes.** Pathway enrichment analysis was performed using Reactome overrepresentation analysis (ORA, collapsed) for intermediate monocytes. Significantly enriched pathways are shown in the left panels and ranked by −log10 adjusted p-value. Bubble size reflects −log10 adjusted p-value, and numbers inside bubbles indicate the number of overlapping genes. Selected genes with high $\Delta\log_2$FC values are shown as trajectory line plots (right), with non-chronic patients in blue and chronic patients in red. Δlog values are indicated per gene panel. Pathway enrichment was performed using Reactome ORA on genes with $|\Delta\log_2 FC| \geq 1$ against a background of all genes tested in the scRNA sequencing differential expression analysis (minSize = 5, maxSize = 500, FDR $\leq$ 0.05). Redundant pathways were removed by iteratively discarding any pathway with a Jaccard similarity $\geq$ 0.60 or smaller-set overlap $\geq$ 0.80 with a higher-ranked pathway, applied to the top 40 significant terms per cell type. Pathogen-specific Reactome terms unrelated to CHIKV were excluded prior to analysis.

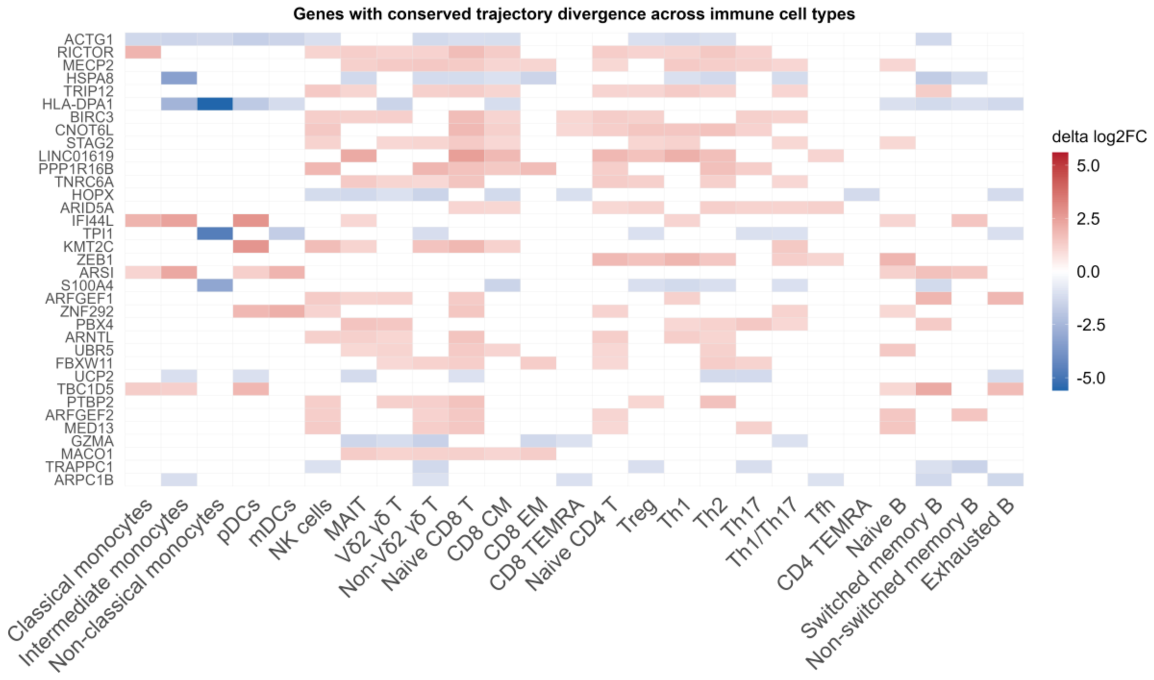


**Supplementary Figure 10. Genes with conserved divergence in longitudinal recovery trajectories across immune cell types.**

Heatmap showing genes with divergent longitudinal expression trajectories between chronic and non-chronic chikungunya patients across immune cell subsets. $\Delta log_2FC$ was calculated as the difference between the acute-to-6-month $log_2$ fold change in chronic and non-chronic patients. Only genes significantly differentially expressed in at least one longitudinal comparison and showing $|\Delta log_2FC| \geq 1$ in $\geq 6$ immune cell types are shown. Color intensity reflects the magnitude of trajectory divergence between groups. Blank cells indicate that the $\Delta log_2FC$ threshold was not met in that cell type.

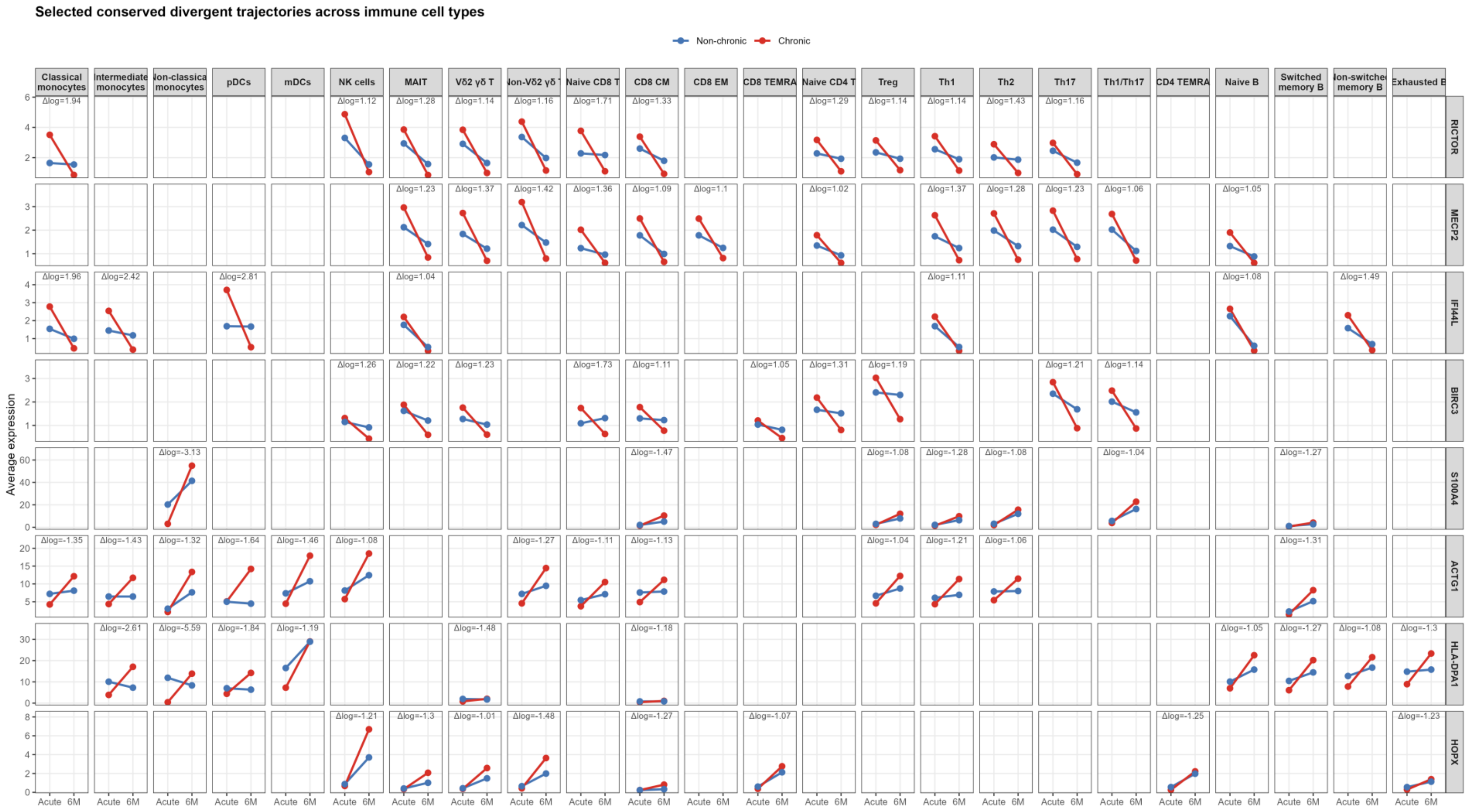


**Supplementary Figure 11: Conserved divergent longitudinal gene-expression trajectories across immune cell subsets.** Line plots showing selected genes with conserved divergent acute-to-6-month expression trajectories across annotated immune cell subsets. For each gene–cell type combination, mean expression is shown at acute infection and 6-month follow-up in non-chronic patients (blue) and chronic patients (red). Panels are faceted by immune cell subset and gene, and only selected conserved trajectory patterns are displayed.

**Supplemental Table 1: Flow cytometry used cell markers**

| Marker | Fluorochrome | Clone | Cat nr | Supplier | |
|---|---|---|---|---|---|
| CD3 | BUV395 | SK7 | 564001 | BD Biosciences | Membrane staining |
| CD4 | BUV496 | SK3 | 612936 | BD Biosciences | Membrane staining |
| CCR7 | BUV737 | 2-L1-A | 749676 | BD Biosciences | Membrane staining 37°C |
| IkBa | BV421 | 3D6C02 | 662409 | Biolegend | Intracellular staining |
| CCR4 | BV605 | L291H4 | 359418 | Biolegend | Membrane staining 37°C |
| CD16 | BV650 | 3G8 | 302042 | Biolegend | Membrane staining |
| CXCR3 | BV711 | G025H7 | 353732 | Biolegend | Membrane staining 37°C |
| CD69 | BV785 | FN50 | 310932 | Biolegend | Membrane staining |
| CCR6 | BB515 | 11A9 | 564479 | BD Biosciences | Membrane staining 37°C |
| CD56 | PE-dazzle | QA17A16 | 392410 | Biolegend | Membrane staining |
| gd TCR | BB700 | 11F2 | 745944 | BD Biosciences | Membrane staining |
| CD45RA | PE cy7 | HI100 | 304126 | Biolegend | Membrane staining |
| TGFBR2 | APC | W17055E | 399706 | Biolegend | Membrane staining |
| CD8 | AF700 | SK1 | 344724 | Biolegend | Membrane staining |
| CD14 | APC-Cy7 | HCD14 | 325620 | Biolegend | Membrane staining |
